# A Phylogenetic Independent Contrast Method under the Ornstein-Uhlenbeck Model and its Applications in Correlated Evolution

**DOI:** 10.1101/2021.04.24.441286

**Authors:** Cong Liang, Yingjun Deng

## Abstract

Phylogenetic comparative methods are essential in studying the evolution of traits across a phylogeny. Felsenstein’s phylogenetic independent contrast (PIC) method and the generalized least squares (GLS) regression were often utilized to study whether evolutionary changes between traits were correlated. However, a neutral Brownian model is assumed in the PIC method, which impacts the performance of the PIC method when the trait is subject to adaptation. In recent years, the Ornstein-Uhlenbeck (OU) model has attracted increasing attention in studying the evolution of traits with stabilizing selection. In this study, we extended Felsenstein’s PIC method under the OU model, which we termed OU-PIC. We simulated trait evolution under the OU model on phylogenetic trees with 8, 10, and 55 species. Compared to the PIC method, the OU-PIC method with correct stabilizing selection parameters achieved an appropriate type I error rate, the highest test power, and the lowest mean squared error. We presented a concise proof of the intrinsic connection between the OU-PIC and the generalized least squares (GLS) regression method in evaluating correlated evolution under the OU model. The OU-PIC method has a broad range of applications when trait evolution could be sufficiently modeled by the OU process. Compared with other phylogenetic comparative methods, OU-PIC avoids the inverse of the covariance matrix and would facilitate the analysis of correlated evolution on large phylogenies.

## Introduction

Correlated evolution studies the relationships between traits or between traits and their environments (Mark Pagel, 1999). By investigating correlated patterns in evolutionary changes, probable causal pathways of trait evolution might be inferred, for example in Podos (2001) and Böhmer et al. (2019). In these studies, traits were measured in species from clades of interest. But species cannot be regarded as independent samples in most instances because their evolutionary histories overlap hierarchically. Therefore, phylogenetically based methods are essential in analyzing such comparative data to ensure the accuracy and reliability of relevant hypothesis testing and inferences (Rohlf, 2006). Felsenstein’s phylogenetic independent contrast (PIC) method (Felsenstein, 1985) has been widely utilized to correct the non-independence of species. The PIC method requires knowledge of the phylogeny with branch lengths and assumes a Brownian motion (BM) model of evolution. The phylogenetic independent contrasts returned by the method (also referred to as PICs, or contrasts) are independent of each other under the BM assumption, and hence are considered suitable for further statistical analysis. Another frequently adopted method for correlated evolution is the generalized least squares (GLS) regression (Martins and Hansen, 1997). The prediction power of one trait on another is evaluated in the GLS regression model. The covariance structure of the error terms describes the relationship among species, and is determined by the phylogeny and the evolution model. The BM assumption is also frequently adopted in the GLS method. Under this assumption, the PIC and the GLS method are intrinsically connected. The slope parameter of the ordinary least squares regression of PIC conducted through the origin is equivalent to the slope parameter of the GLS parameter in evaluating correlated evolution (Rohlf, 2006; Blomberg et al., 2012).

The Brownian model of evolution is neutral and has no adaptation term. In practice, the BM assumption of trait evolution is likely violated. Multiple simulation studies have evaluated the performance of different phylogenetic comparative methods (PCMs) when the underlying evolutionary process deviates from BM (Martins and Garland, 1991; Martins et al., 2002). Among these methods, the PIC method has been shown to outperform others in most cases and is generally acceptable in studying correlated evolution even if the underlying evolution model is not BM (Martins and Garland, 1991). A widely utilized alternative to the BM model is the Ornstein-Uhlenbeck (OU) model, where both the effect of stabilizing selection as well as random drift were modeled (Hansen, 1997; Butler and King, 2004). In contrast to the BM model where the variance of traits increases without limit, the trait value tends to drift around its optimal level with a bounded variance under the OU model. This bounded property made OU the preferred model in many evolutionary scenarios, for example in describing the evolution of molecular traits like gene expression (Bedford and Hartl, 2009; Yang et al., 2019). However, the covariance structure of species is more complicated under the OU model (Hansen and Martins, 1996; Hansen, 1997). More and more analytical methods were established to study trait evolution under the OU model in recent years (Rohlfs et al., 2014; Khabbazian et al., 2016). Difficulties have also been addressed in studying evolutionary questions with OU models (Ho and Ané, 2013, 2014).

In this manuscript, we derived a generalization of Felsenstein’s PIC method under the OU model of trait evolution, which we termed as the OU-PIC method. Our method calculates *n* − 1 phylogenetic independent contrasts (OU-PICs) from observations in *n* species according to their phylogenetic structure. We modeled the level of correlated evolution by the correlation coefficient between the stochastic terms of OU processes, which enabled us to construct explicit statistical tests and inferences on correlated evolution. We studied the correlated evolution of continuous traits under the OU model using the OU-PIC method. Our simulation results showed that OU-PIC improves the statistical performance of the PIC method under the OU model. When the correct model parameter is specified, OU-PIC delivers appropriate type I error rates, obtained the highest power of tests, and the lowest mean squared errors in estimations. Finally, we highlighted the intrinsic connection between the OU-PIC and the GLS regression method by analyzing the contrast coefficient matrix.

## Materials and Methods

### The Ornstein-Uhlenbeck Model of Trait Evolution

We consider the Ornstein-Uhlenbeck model for the evolution of a continuous trait *X* (Hansen, 1997; Butler and King, 2004). The evolution process is described by the following stochastic differential equation,

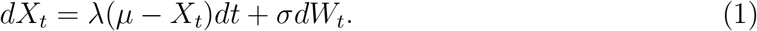

*W*_*t*_ is a Wiener process or standard Brownian motion. It has Gaussian increments and its variance is linear to time *t*. The stochastic term, *σdW*_*t*_, models the effect of random drift with strength parameter *σ*. The deterministic term, λ(*μ* − *X*_*t*_)*dt*, models the effect of stabilizing selection. Parameter λ measures the strength of stabilizing selection and is always positive. With stabilizing selection, *X*_*t*_ tends to drift towards its fitness optimum *μ*. If *X*_*t*_ deviates far from *μ*, the character value will be pulled towards its optimum with a large slope. If λ = 0, Equation (1) becomes the Brownian model.

Equation (1) has analytical solutions, which makes it a tractable model for studying trait evolution. Given an ancestral state *x*_0_, *X*_*t*_ follows a normal distribution, with known expectation and variance.

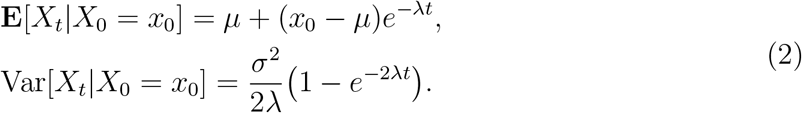

The expectation of *X*_*t*_ decays or increases exponentially with time.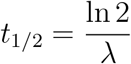, also called the half-time, is when **E**[*X*_*t*_] is halfway towards its optimal. For a trait evolving along a known phylogeny, the proportion of the half-time to the time span of the phylogeny describes how fast stabilizing selection pulls *X*_*t*_ to its optimal relative to the time span of the phylogeny. Take *L* as the distance from the root to current species on the phylogeny. We define κ as the relative strength of stabilizing selection, which is the ratio of *L* and *t*_1/2_,

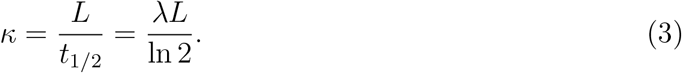

If *X* has evolved a very long time *τ*, *X*_*τ*_ follows a normal distribution that is independent of *x*_0_ and *τ*,

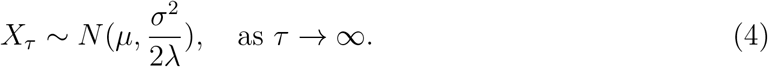

### Trait Evolution along a Phylogenetic Tree Under the OU Model

To study the evolution of a trait *X* along a known phylogeny (for example as in Figure 1), the time interval between the last common ancestor of all species (*t* = 0) and current time (*t* = *t*_0_) is generally considered. In fact, most traits, especially molecular traits such as gene expression levels, could only be observed at the current time, i.e., *t* = *t*_0_. As a result, we drop the time index of trait values and denote its value on tip node *k* of the phylogeny by random variable *X*_*k*_; also denote the trait value on the root of the phylogeny by random variable *X*_0_. We made several general assumptions in this study: (1) Branch lengths on the phylogeny are in common clock scale. (2) Only extant species are considered, so all tip nodes obtain the same distance to the root of the tree. (3) The optimal trait value has not changed in the course of evolution. (4) When the evolutionary history overlapped, the evolution of *X*_*k*_ are the same process. They became independent after the speciation event.

**Figure 1.**
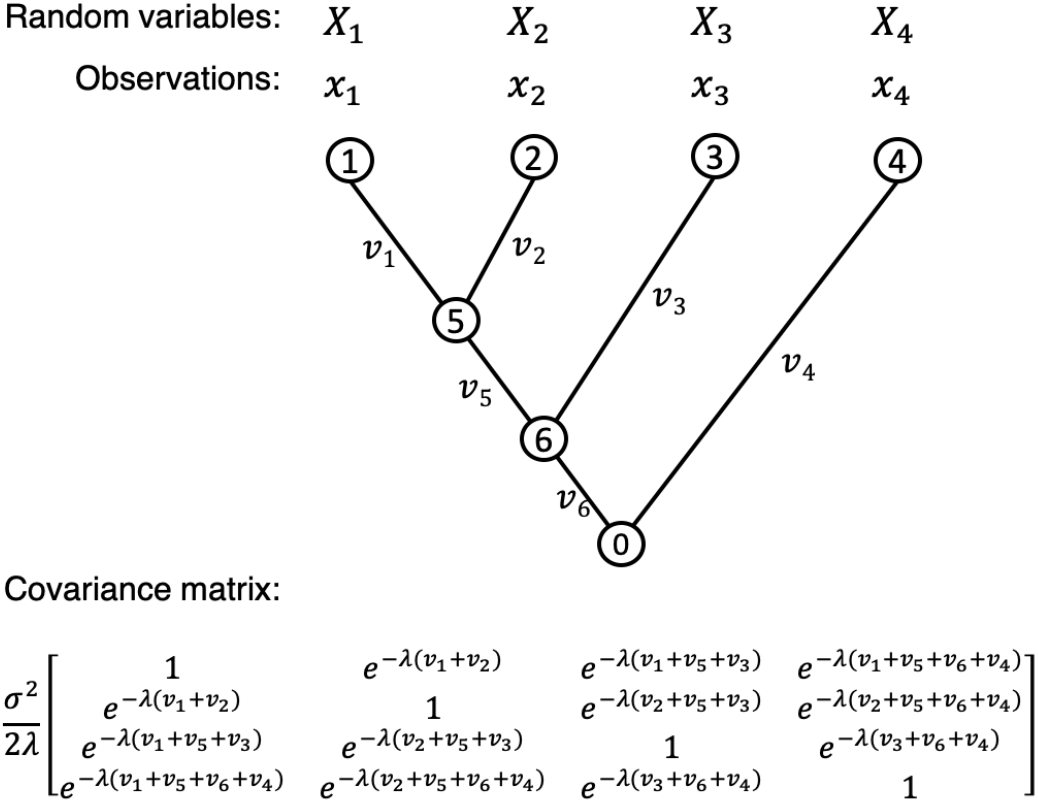
An illustration of a 4-species phylogeny and the evolution of trait *X*. Extant species are indexed from 1 to 4, and the last common ancestor of all species is indexed as node 0. Trait values are only observed in extant species, i.e., tips of the species tree. We used capitalized letters, *X*_1_,… , *X*_4_ for random variables of trait *X* in different species, and the corresponding lower letters, *x*_1_,… , *x*_4_ for their observed values. When their evolutionary history overlapped, the evolution of *X*_*k*_ are the same process, and processes in two species became independent after speciation. The branch lengths are in common clock scale and are given as *ν*_*i*_. The distance from the root to the tips of the species tree is a constant. We illustrated the covariance matrix of ***X*** = (*X*_1_,… , *X*_4_) according to Equation (5).

Further, if trait *X* originated long before the root of the phylogeny, the evolution time *τ* between the origin of *X* and the root of the phylogeny is so long that we take *τ* → ∞. *X*_0_ and all *X*_*k*_ follow the stationary distribution in Equation (4) that 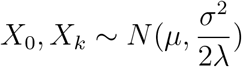. This ancient-origin assumption has been adopted in multiple studies and was also called the stationary assumption of the OU model (Hansen, 1997; Bedford and Hartl, 2009; Yang et al., 2019). We will discuss trait evolution under the OU model with this assumption throughout this manuscript.

As under the Brownian model, trait values on different nodes of the tree are correlated to each other under the OU model. The covariance between two nodes *i* and *j* could be derived using the analytical solution of OU processes (Appendix I, also in Hansen and Martins (1996))

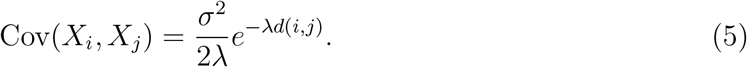

*d*(*i*, *j*) is the divergence time between *i* and *j*. We illustrated a simple phylogenetic tree and the covariance matrix for its tip node values in Figure 1.

### Correlated Evolution under the OU Model

We studied correlated evolution between two traits *X* and *Y* under the OU model (Liang et al., 2018). Both evolutionary trajectories follow the Ornstein-Uhlenbeck process:

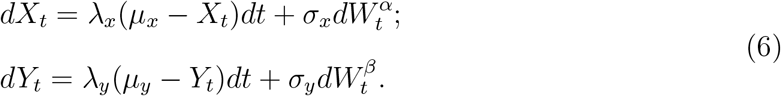

We introduced a parameter *γ*_*xy*_ for the level of correlated evolution between *X* and *Y*. It is the correlation between the two Wiener processes 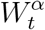 and 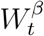:

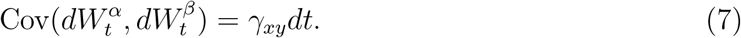

Alternatively, 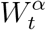 and 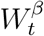 could be considered as linear combinations of two independent Wiener processes 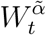 and 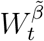

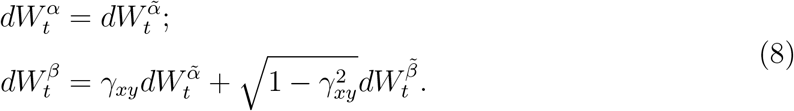

According to this model, *γ*_*xy*_ ϵ (−1, 1). *γ*_*xy*_ = 0 indicates the evolution of *X* and *Y* are independent to each other; whereas *γ*_*xy*_ close to −1 or 1 indicates strong correlation between the evolution of *X* and *Y*. *γ*_*xy*_ = ±1 are mathematically plausible but are biologically unrealistic. To address questions regarding correlated evolution between *X* and *Y*, we investigate hypothesis tests and inferences with respect to parameter *γ*_*xy*_.

We derive the covariance between *X*_*i*_ (trait *X*’s value on node *i*) and *Y*_*j*_ (trait *Y*’s value on node *j*) under the OU model with stationary assumption (Appendix II).

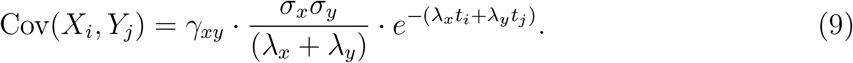

Here, *t*_*i*_, *t*_*j*_ are the evolution time between *i*, *j* and their last common ancestor *k*, respectively. The joint distribution of character values in *n* extant species (*X*_1_,… , *X*_*n*_, *Y*_1_,… , *Y*_*n*_)^*T*^ is multivariate normal with mean (*μ*_*x*_,… , *μ*_*x*_, *μ*_*y*_,… , *μ*_*y*_) and elements in covariance matrix Σ_*xy*_ as in Equation (9).

### Statistical Performance of OU-PICs in Evaluating Correlated Evolution

We estimated the level of correlated evolution *γ*_*xy*_ using the Pearson correlation coefficient between OU-PICs of two traits *X* and *Y*, 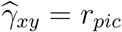 (see Results). According to Appendix III, *r*_*pic*_ is an unbiased estimator of *γ*_*xy*_ when λ_*x*_ = λ_*y*_ (or equivalently, *κ*_*x*_ = *κ*_*y*_). We evaluated the statistical performance of *r*_*pic*_ in estimating *γ*_*xy*_ with respect to type I error rates, the power of test, and the mean squared error.

#### Type I Error Rate

We evaluated the type I error rates of using OU-PICs to test whether there is correlated evolution between two traits. The null hypothesis, *H*_0_: *γ*_*xy*_ = 0. The alternative hypothesis, *H*_1_: *γ*_*xy*_ ≠ 0. The type I error rate is the probability of rejecting the null hypothesis when it is in fact true. We simulated trait values with no correlated evolution (*γ*_*xy*_ = 0) under the OU model of evolution as well as the Brownian model. The level of relative stabilizing selection varies, *κ* ϵ (2^0^, 2^1^, 2^2^, 2^3^, 2^4^, 2^5^). *κ* were set the same for *X* and *Y* in each simulation. Theoretically, when *H*_0_ is true, a transformation of the Pearson correlation coefficient, 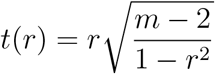 follows Student’s t-distribution with degrees of freedom *df* = *m* − 2. *m* here is the number of samples. When applying the PIC or OU-PIC method, *m* equals to the number of contrasts. For a *n*-species phylogeny, *m* = *n* − 1. When no comparative method was applied (“Raw Tips” in Fig. 5), *m* equals to the number of species. We determined the two-sided critical values *t*_*c*_(*α*) for varied *α* levels with **P**(|*t*| < *t*_*c*_(*α*)) = 1 − *α*, and then identified the corresponding critical values for Pearson correlation coefficients, *r*_*c*_(*α*). The type I error rates were calculated by the fraction of observed *r*_*pic*_ beyond critical values, |*r*_*pic*_| > *r*_c_(*α*).

#### Power of Test

We evaluated the power of the above hypothesis test when *H*_1_ is true. We simulated the evolution of two traits *X* and *Y* with varied levels of correlated evolution, *γ*_*xy*_ ϵ (0, 0.25, 0.5, 0.75, 0.9). The level of stabilizing selection also varies, *κ* ϵ (0, 2^0^, 2^1^.2^2^, 2^3^, 2^4^, 2^5^). The critical values *r*_*c*_(*α*) of the test were determined using the empirical distributions of *r*_*pic*_ when *H*_0_ is true; **P**_*H*_0__ (|*r*_*pic*_| ≤ *r*_*c*_(*α*)) = 1 − *α*. The power of test was then estimated by the fraction of *r*_*pic*_ beyond the critical values when *H*_1_ is true; power = **P**_*H*_1__ (|*r*| > *r*_*c*_(α)).

#### Mean Squared Error

We calculated the mean squared error in estimating *γ*_*xy*_ using *r*_*pic*_ with *MSE* = Σ(*r*_*pic*_ − *γ*_*xy*_)^2^.

### Phylogenetic Trees Analyzed in This Study

We tested OU-PICs on phylogenetic trees with a varied number of species and structures as illustrated in Figure 2. (1) An 8-species tree of Darwin’s finches (Finch Tree). This is one of the four phylogenetic hypotheses adopted in Podos (2001) to study the correlated evolution of morphology and vocal signal structures in Darwin’s finches. Branch lengths in this tree are arbitrary, and not necessarily in time scales. We chose this tree as a representation of small and relatively balanced phylogenies. (2) A 10-species tree containing 9 mammalian species and one bird (Brawand Tree). This is the species sampling tree in Brawand et al. (2011) to study the evolution of gene expression in mammalian organs. The study sampled broadly among mammals but also has a special focus on great apes. We chose this tree because Brawand et al. (2011) is one of the most frequently utilized resources to study transcriptome evolution, plus this tree has a special comb-like structure with highly varied branch lengths. (3) A 55-species tree of maples (genus Acer, Acer Tree). The correlated evolution of leaf size, sapling canopy allometry, and other related traits of Acer species have been studied in Ackerly and Donoghue (1998). We chose this tree to represent larger phylogenies. The time tree of Acer species were downloaded from the time tree of life project (http://www.timetree.org/) (Hedges et al., 2015). Ambiguous nodes were removed manually from this tree in this study. We note that our OU-PIC method can also handle hard polytomy by setting some branch lengths as zero. We simulated values of two traits in extant species on these trees by direct sampling from their joint multivariate distributions. The number of simulations *N* = 10, 000.

**Figure 2.**
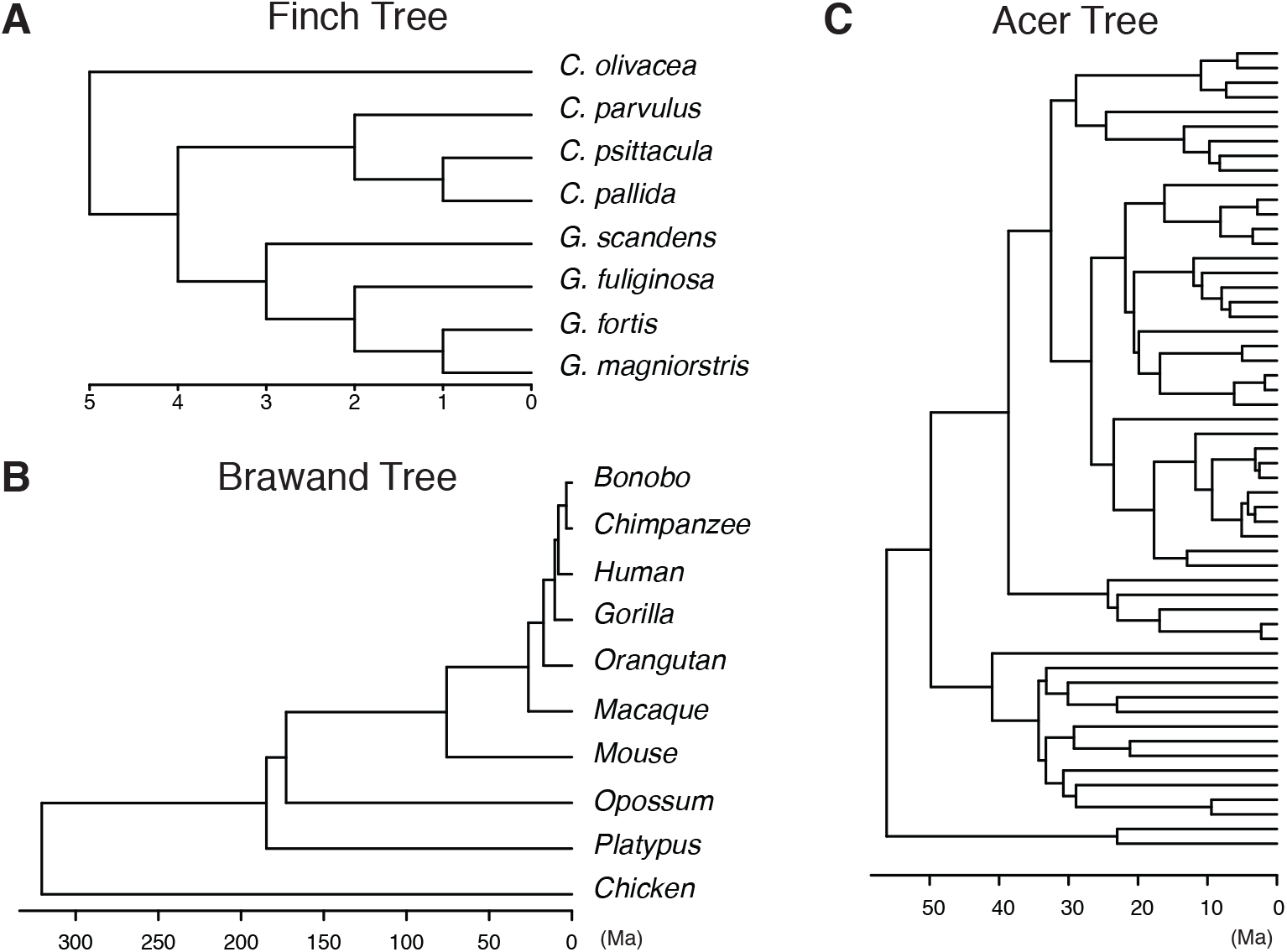
Phylogenetic trees analyzed in this study. (A) An 8-species tree of Darwin’s finches studied in Podos (2001) (Finch Tree). The branch lengths are arbitrary. (B) A 10-species tree containing nine mammals and one brid (Brawand Tree). This tree was sampled in Brawand et al. (2011) to study the evolution of gene expression in mammals with a focus on great apes. This tree has a comb-like structure, and its branch lengths vary heavily. (C) A 55-species tree of Acer species studied in Ackerly and Donoghue (1998) (Acer Tree). Ambiguous nodes were manually removed from this tree in this analysis. Speciation times in (B) and (C) are extracted from the time tree of life project (http://www.timetree.org/) (Hedges et al., 2015).

## Results

### A Phylogenetic Independent Contrast Method for the OU model

We adopted a tree pruning procedure analogous to Felsenstein’s PIC method to calculate phylogenetic independent contrasts under the OU model (Algorithm 1). The algorithm traverses the internal nodes of a species tree according to the post-order and prunes their descendants recursively (Fig. 3). The post-order traversal is often used to delete a tree since the ancestor node is visited after its descendants in this procedure. In each iteration of Algorithm 1, two descendants of an internal node **w**_*i*_ were removed, making **w**_*i*_ a new leaf of the pruned tree. We denoted the two descendants of **w**_*i*_ by x_*l*_ and x_*r*_, let *x*_*l*_ and *x*_*r*_ denote their trait values, and let *ν*_*l*_, *ν*_*r*_ denote their distances to **w**_*i*_. Then, we calculated a contrast *c*_*i*_, which is a weighted difference between *x*_*l*_ and *x*_*r*_, and an intermediate value *w*_*i*_, which is a weighted average of *x*_*l*_ and *x*_*r*_.

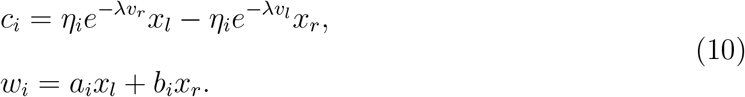

**Figure 3.**
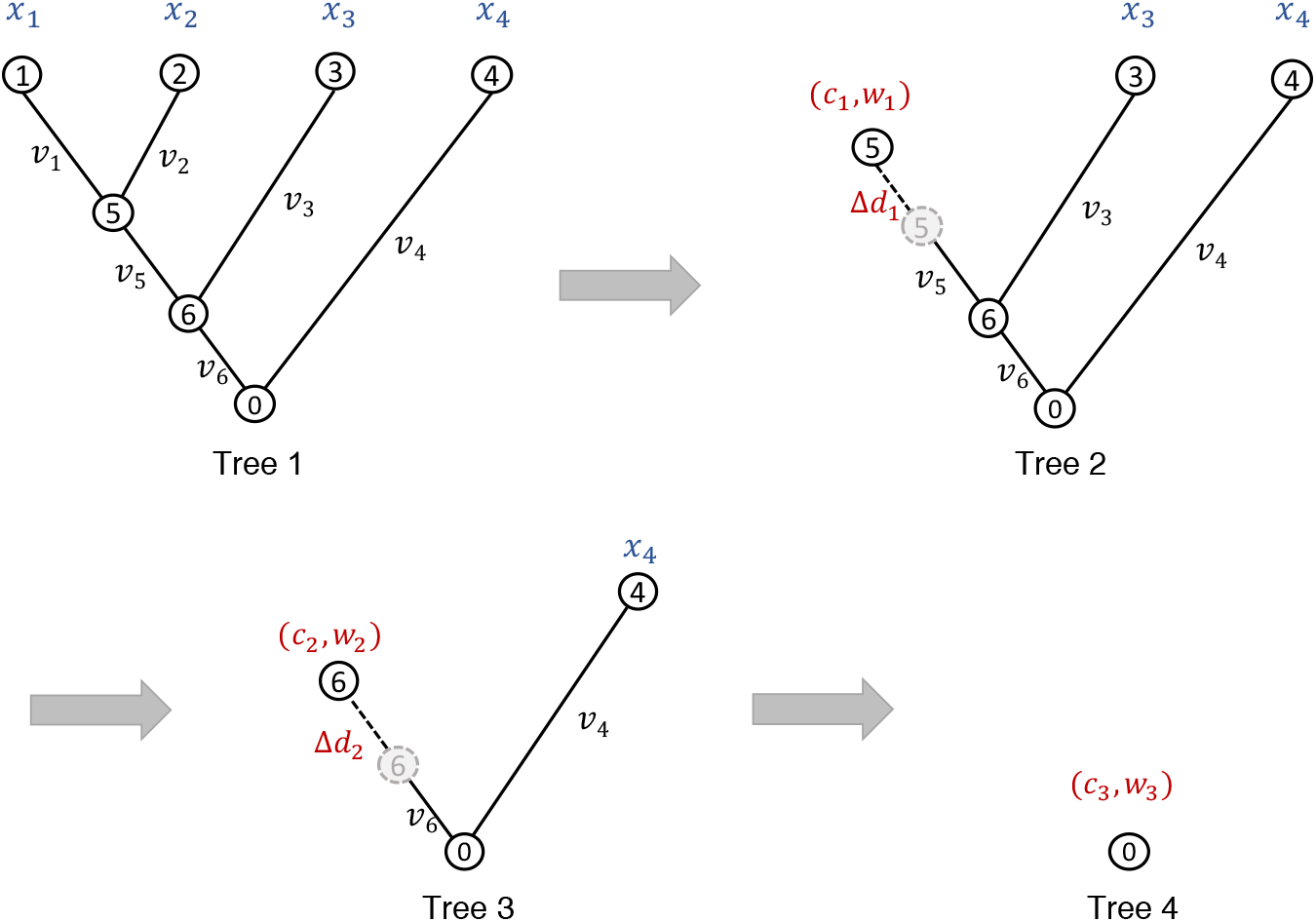
An illustration of the tree pruning process to calculate OU-PICs following Algorithm 1. The procedure is analogous to the tree pruning procedure in Felsenstein’s PIC method. Internal nodes were visited by the order 5 → 6 → 0. In each iteration, two leaf nodes were trimmed and their last common ancestor became a leaf in the pruned tree. A contrast *c*_*i*_ and an intermediate node value *w*_*i*_ were calculated in each iteration. *c*_*i*_ is returned as the *i*-th contrast by the algorithm. *w*_*i*_ is assigned to the new leaf node in the pruned tree. A modification of branch length Δ*d*_*i*_ is calculated between the new leaf node and its nearest ancestor. The process continues until only one node remained. A total of *n* − 1 contrasts and intermediate values were calculated from this procedure.

Here, e^λ*ν*_*r*_^ and −e^λ*ν*_*l*_^ make the mean value of the contrast zero. *η_i_* is the normalizing factor that makes the variances of all contrasts consistent. *a*_*i*_ and *b*_*i*_ were determined by the condition that the covariance between the contrast and the intermediate value is zero, and that intermediate values have consistent covariances. *c*_*i*_ is the *i*-th contrast returned by the algorithm. *w*_*i*_ is assigned to node **w**_*i*_ in the pruned tree. At the end of each iteration, we modified the branch length between **w**_*i*_ and its nearest ancestor by 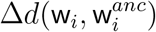 to make the covariance matrix among leaf nodes maintain the same structure in the pruned tree. These processes were iterated until there is only one node left on the pruned tree.

We specified the formulas of *η*_*i*_, *a*_*i*_, *b*_*i*_ and 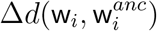 in Equation (23) (Appendix III). Our algorithm assumed that the stabilizing selection parameter λ is known along with the species tree. In practice, λ could be estimated using maximum likelihood or other methods designed for the relevant data type (Bedford and Hartl, 2009; Gu et al., 2019).

Our algorithm returns *n* − 1 contrasts (OU-PICs) for traits observed in n extant species. To describe the statistical characteristics of our method, we distinguish random variables from their observed values in notations. We use capitalized letters to indicate random variables and the corresponding lowercase letters for observed or calculated values. For example, *C*_*i*_ stands for an independent contrast random variable, whereas *c*_*i*_ represents a contrast calculated from observed trait values. We showed that the OU-PICs have the following properties (proof in Appendix III):

1. *C*_*i*_ are independent and identically distributed following a normal distribution; *C*_*i*_ ~ *N*(0, *ϕ*), the normalized variance 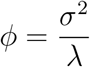
2. The contrasts of two traits *X* and *Y* are pairwise correlated and their correlation equals the correlated evolution parameter *γ*_*xy*_, if they obtain the same level of stabilizing selection, λ_*x*_ = λ_*y*_.

Like Felsenstein’s PIC method, our OU-PIC method removes the correlation among species resulted from overlapping evolutionary histories. For two traits *X* and *Y* with the same level of stabilizing selection on a known phylogeny, the Pearson correlation coefficient between their OU-PICs is an unbiased estimator of their level of correlated evolution *γ*_*xy*_.

#### Algorithm 1

Phylogenetic Independent Contrast Method under the OU Model

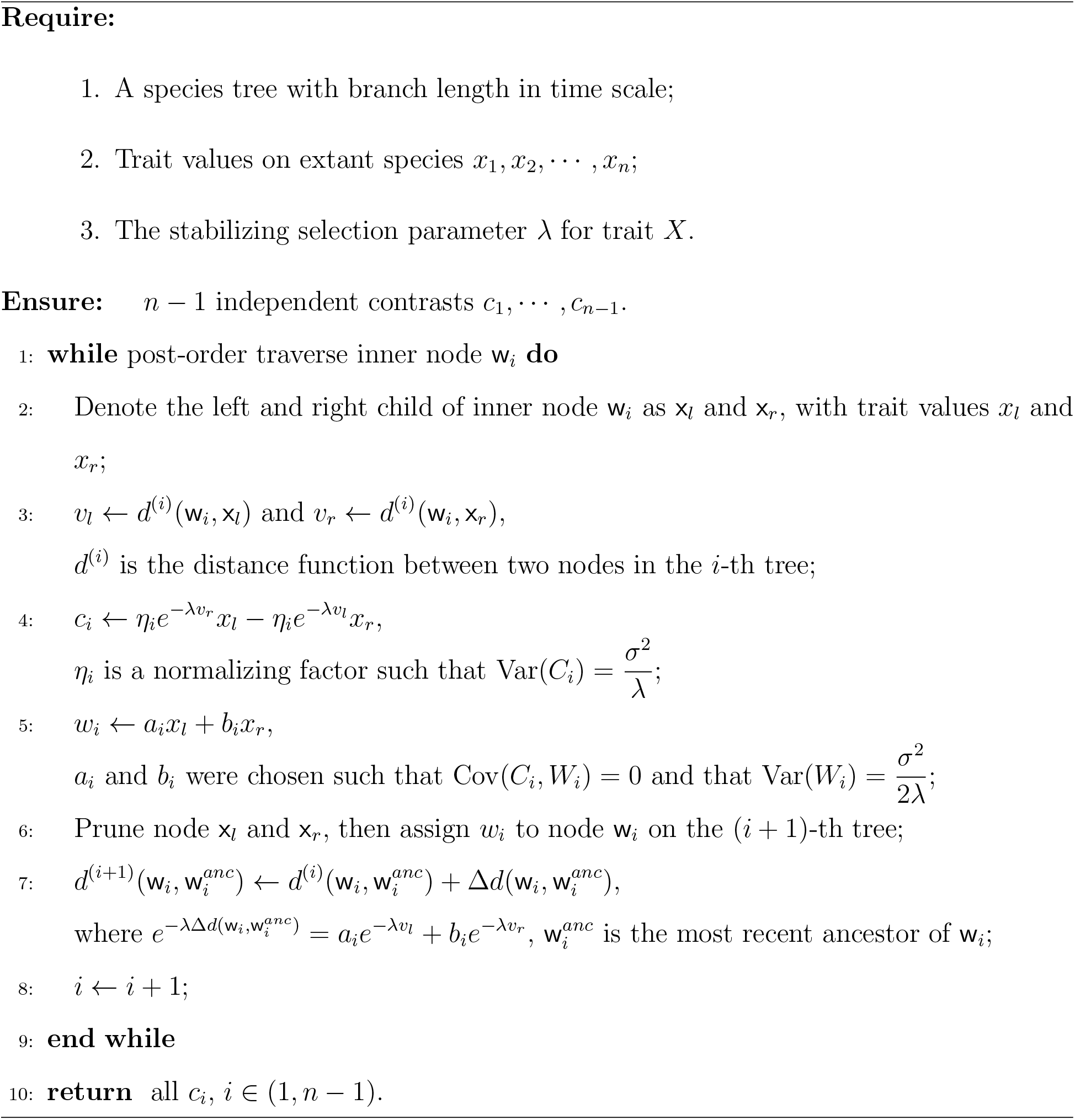

### The Coefficient Matrix of OU-PICs

According to Algorithm 1, OU-PICs are linear combinations of observed trait values. We rewrote OU-PICs as 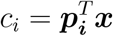. Here, ***x*** = (*x*_1_,… , *x*_*n*_)^*T*^ are observed trait values in extant species. ***p***_i_ (*i* = 1,… , *n* − 1) are contrast coefficient vectors each with *n* elements. Bold faces denote column vectors throughout this paper. Let ***c*** = (*c*_1_,… , *c*_*n*-1_)^*T*^ denote the contrast vector. We have,

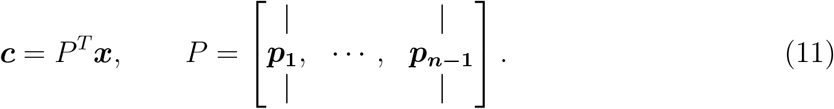

We call *P* the contrast coefficient matrix. Knowing that the contrasts are independent identically distributed random variables, the covariance matrix of the contrast vector is diagonalized. We have,

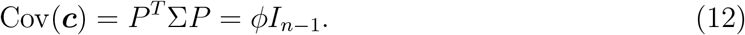

Here, *I*_*n*-1_ is an order *n* − 1 identity matrix. In fact, Equation (12) is also true for Felsenstein’s PICs. Both PICs and OU-PICs are linear transformations of trait values such that the covariance matrix is diagonalized under given evolutionary assumptions. The OU-PIC method is a generalization of Felsenstein’s PIC method to OU assumptions.

The contrast coefficient matrix of OU-PICs differs from that of PICs in its non-zero terms. Under the BM model, accumulated variances in traits increase linearly with evolution time without limit. Whereas under the OU model, the variances of traits are constrained by stabilizing selection. Therefore, contrasts associated with long branches tend to be normalized with a greater variance in the PIC method than in the OU-PIC method. To compare the strength of stabilizing selection across species trees, we defined a dimension one quantity, *κ*, which describes how fast stabilizing selection drags the trait value towards its optimal relative to the time span of the phylogeny. Specifically, *κ* is proportional to the product of λ and the distance between the root and tips of the species tree (see Methods).

We observed shrunken coefficients of PICs associated with long branches compared to that of OU-PICs (Fig. 4 and Supp. Fig. 1). When the relative stabilizing selection is moderate or strong, for instance *κ* > 10 for the Finch tree and the Brawand tree, the coefficients of OU-PICs deviate from the coefficients of PICs substantially. As *κ* decreases, the differences between the coefficients of OU-PICs and the coefficients of PICs decline. When the relative stabilizing selection is weak, for instance *κ* ≤ 1 for the Finch tree and the Brawand tree, the coefficients of OU-PICs are close to that of PICs. In this case, the evolution process of traits on the species tree can be approximated by a BM process. We also observed that the differences between the coefficients of OU-PICs and PICs are more significant in the Brawand tree than in the Finch tree, because branch lengths vary more heavily in the Brawand tree.

**Figure 4.**
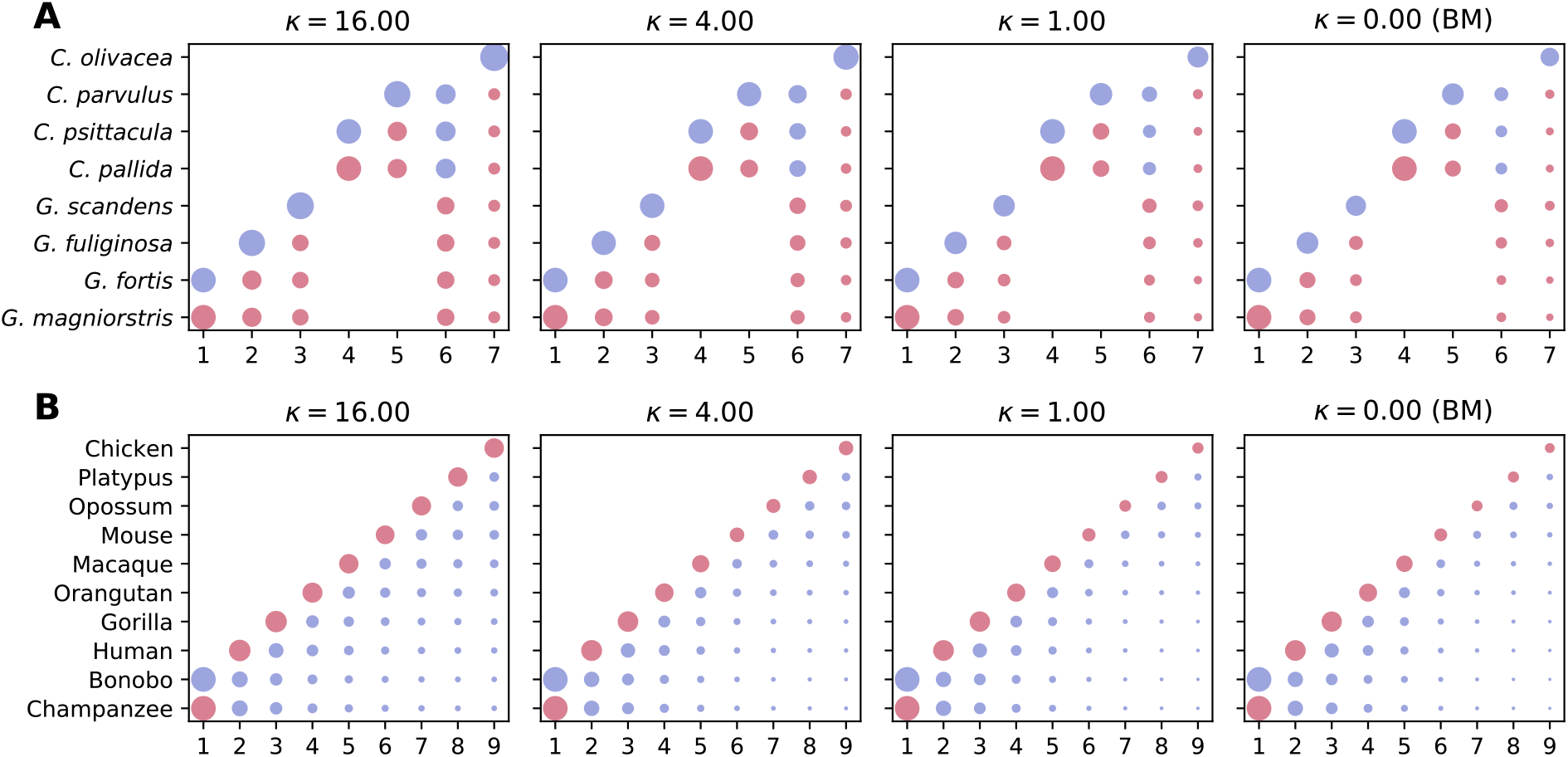
Visualization of the coefficient matrix of OU-PICs and PICs for (A) the Finch Tree, and (B) the Brawand Tree. The level of relative stabilizing selection varies under the OU model. When *κ* = 0, the evolution model is BM, and the coefficient matrix of PICs is plotted. The horizontal axis in each panel represents the index of contrasts. The vertical axis in each panel represents extant species on a species tree. To compare coefficients for different *κ*, coefficients were normalized according to the coefficients of contrast 1 in each panel (the bottom left value). The size of the points is proportional to the absolute value of the normalized coefficients. The color of points represents the sign of coefficients, red for positive, blue for negative. We observed distinct coefficients for PICs and OU-PICs in contrasts associated with long branches, for example contrasts 3 and 7 in the Finch Tree, and contrasts 7-9 in the Brawand Tree. The difference between coefficients of PICs and OU-PICs increases with *κ*.

### Evaluating Correlated Evolution Using OU-PICs

To evaluate the statistical performance of OU-PICs in studying correlated evolution, we simulated the evolution of two traits *X* and *Y* with varied levels of correlated evolution and stabilizing selection. The strength of stabilizing selection for two traits were set equal in the simulation, *κ*_*x*_ = *κ*_*y*_ = *κ*. The contrasts were calculated using varied assumed levels of stabilizing selection 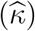. when 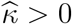, the OU-PIC method was applied to get the contrasts. When 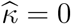, the PIC method is utilized. The level of correlated evolution was then estimated by the Pearson correlation coefficient between the contrasts of two traits, 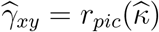. We compared the statistical performance of the PIC and the OU-PIC method with respect to (1) the type I error rate when *γ*_*xy*_ = 0; (2) the power of test when *γ*_*xy*_ > 0; and (3) the mean squared error in estimating *γ*_*xy*_ (see Methods).

We found that OU-PIC achieved appropriate type I error rates when 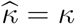, i.e. the correct level of stabilizing selection was assumed in the analysis (Fig. 5). If 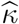 deviates from *κ* by only several folds, the type I error was slightly inflated. For example, the type I error is always smaller than 0.06 for all three species trees when the difference between 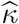 and *κ* is less than two folds and *α* = 0.05. However, if 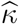 deviates excessively from *κ*, type I error rates could be highly inflated. Notably, the type I error rate was inflated by more than two folds if the PIC is inappropriately adopted when the underlying stabilizing selection is strong. For example, the type I error = 0.136 for the Brawand tree and 0.173 for the Acer tree when *κ* = 32, 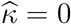 and α = 0.05. Similarly, we obtained the highest power of test (Fig. 6 and Supp. Fig. 2) and the smallest mean squared error in estimating *γ*_*xy*_ (Supp. Fig. 3 and 4) when the appropriate *κ* is specified 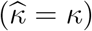. When comparing species trees, we found that the Brawand tree and the Acer tree are more sensitive to parameters and model assumptions than the smaller Finch tree. Overall, these results suggest that it is important to evaluate the impact of stabilizing selection in comparative studies. Methods should be adopted accordingly to achieve the desired statistical performance when evaluating correlated evolution. When *κ* is small (*κ* ≪ 1), the BM model is sufficient to describe the underlying evolutionary process. However, as *κ* increases (*κ* > 1), the BM model can no longer serve as an adequate model. The statistical performance of the PIC method drops significantly. Specifically designed methods such as the OU-PIC method could improve the statistical performance under such circumstances.

**Figure 5.**
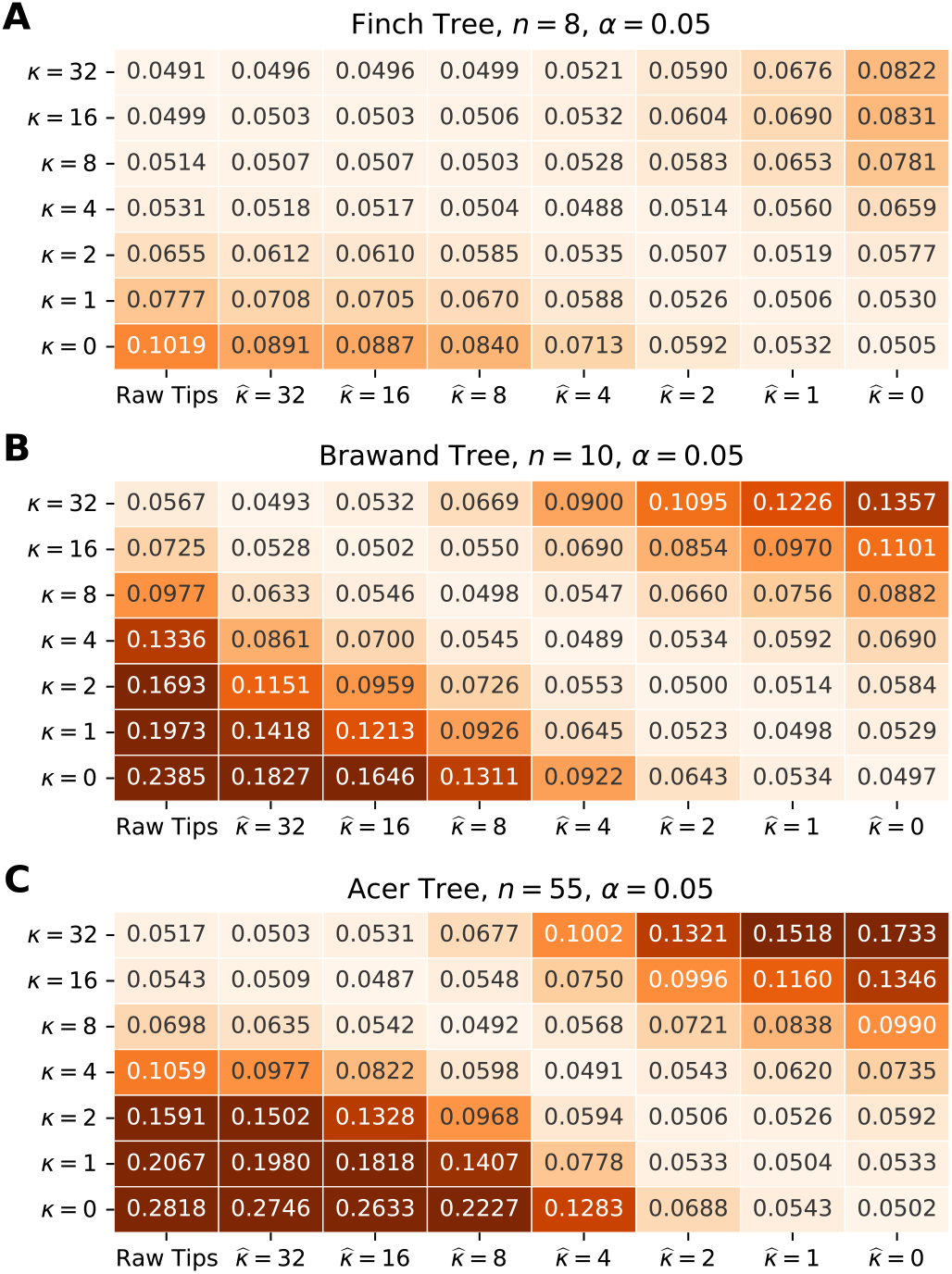
Type I error rates in testing whether there is correlated evolution on (A) the Finch Tree, (B) the Brawand Tree, and (C) the Acer Tree. We simulated the evolution of two traits with no correlated evolution, and estimated the fraction of 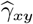 beyond the theoretical threshold for α = 0.05. The vertical axis in each panel is the true level of stabilizing selection in simulation, *κ*. The horizontal axis in each panel is the level of stabilizing selection used for calculating OU-PIC. The first column, “Raw Tips”, correlated evolution is estimated with no comparative method, i.e. the correlation between observed values in extant species. When *κ* = 0, the true evolution model is BM. When 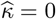, PICs were utilized in the analysis. We observed appropriate type I errors when the correct level of correlated evolution is specified in the analysis, 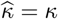 (diagonal of the table). Type I error rates were inflated when 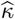 deviate from *κ*.

**Figure 6.**
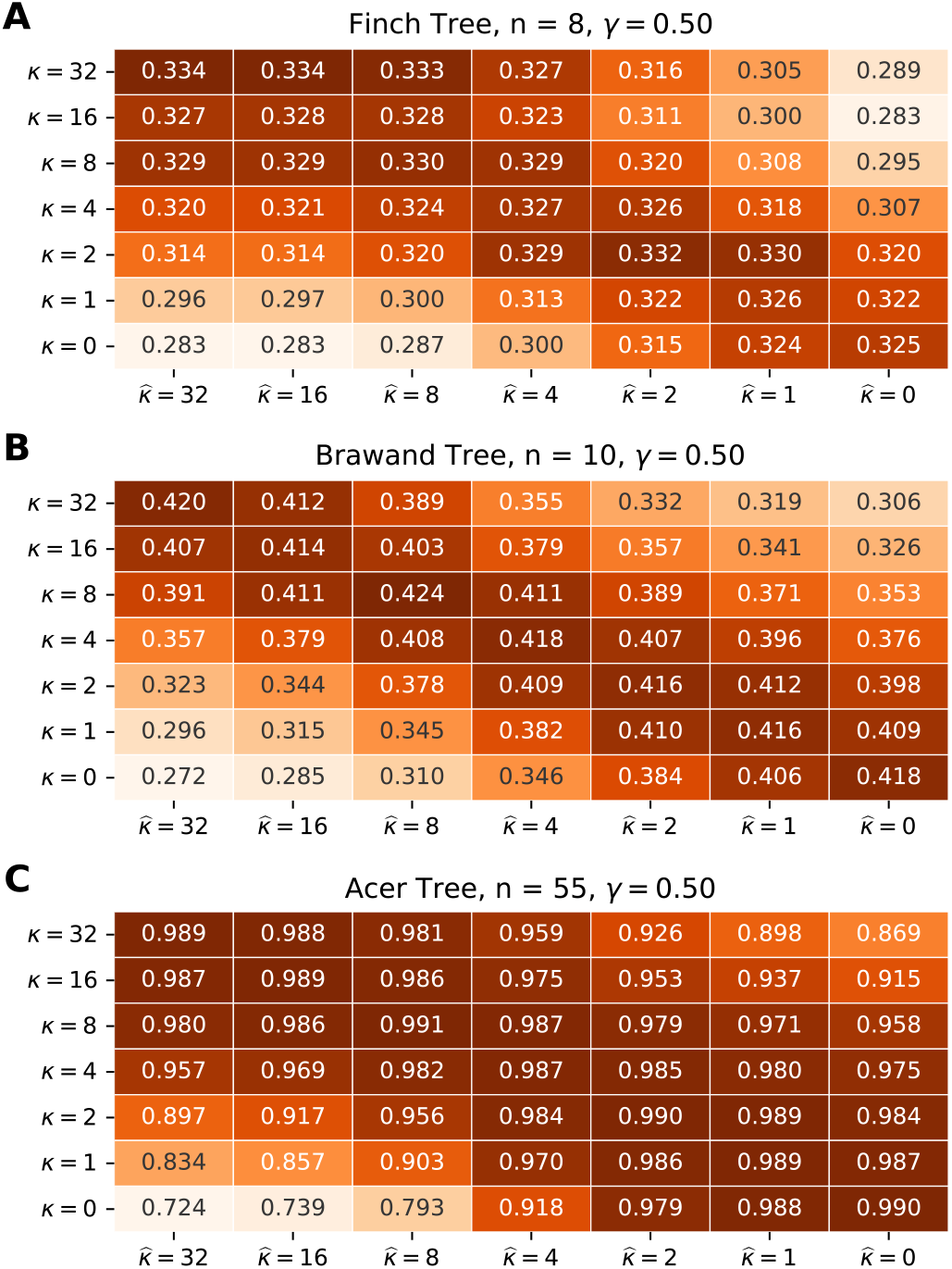
The power of test on (A) the Finch Tree, (B) the Brawand Tree, and (C) the Acer Tree given, *γ* = 0.5. The highest test power is achieved when the correct level of stabilizing selection is specified, i.e. 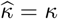.

### Correlated Evolution with Unequal Stabilizing Selection Parameters

When *κ*_*x*_ ≠ *κ*_*y*_, the covariance matrix of OU-PICs for *X* and *Y* becomes analytically complicated. Pairs of OU-PICs, 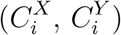 for i = 1, 2,… , *n* − 1, become inter-correlated in this case. As a result, we tested the performance of OU-PICs in evaluating correlated evolution when *κ*_*x*_ ≠ *κ*_*y*_ numerically. In practice, if *κ*_*x*_ and *κ*_*y*_ dramatically differ from each other (e.g. by 5 folds or more), the effect of stabilizing selection on their evolution processes also diverges significantly. It does not make much biological sense to study correlated evolution between such two traits. Therefore, we simulated trait evolution with slightly different *κ*_*x*_ and *κ*_*y*_, (*κ*_*x*_, *κ*_*y*_) = (4.5, 5.5), (3, 7) or (2, 8). We again evaluated the type I error rates, the power of tests, and mean squared errors in studying the level of correlated evolution using the OU-PIC method (assuming 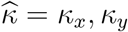 or (*κ*_*x*_ + *κ*_*y*_)/2) and the PIC method 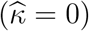. We found that in all three phylogenetic trees we analyzed, the OU-PIC method still obtained appropriate statistical performance and outperformed the PIC method (Table 1).

**Table 1.** The type I error rates, test power, and mean squared errors using the PIC and OU-PIC method to evaluate correlated evolution when *κ*_*x*_ ≠ *κ*_*y*_.

| | $\kappa_x$ | $\kappa_y$ | $\hat{\kappa}$ | Type I error rate | $\gamma = 0.25$ | | | | $\gamma = 0.50$ | | | | $\gamma = 0.75$ | | | |
| --- | --- | --- | --- | --- | --- | --- | --- | --- | --- | --- | --- | --- | --- | --- | --- | --- |
|  |  |  |  |  | power | mean | std | mse | power | mean | std | mse | power | mean | std | mse |
| Finch Tree (n=8) | 4.5 | 5.5 | 4.5 | 0.051 | 0.133 | 0.230 | 0.389 | 0.152 | 0.328 | 0.466 | 0.337 | 0.115 | 0.696 | 0.715 | 0.230 | 0.054 |
|  |  |  | 5.5 | 0.049 | 0.133 | 0.231 | 0.390 | 0.153 | 0.322 | 0.465 | 0.336 | 0.114 | 0.697 | 0.716 | 0.230 | 0.054 |
|  |  |  | 5.0 | 0.049 | 0.137 | 0.231 | 0.390 | 0.153 | 0.330 | 0.466 | 0.337 | 0.115 | 0.700 | 0.716 | 0.228 | 0.053 |
|  |  |  | 0 (PIC) | **0.069** | *0.127* | *0.227* | *0.417* | **0.175** | *0.297* | *0.462* | *0.361* | **0.132** | *0.637* | *0.712* | *0.248* | **0.063** |
|  | 4.0 | 6.0 | 4.0 | 0.051 | 0.133 | 0.228 | 0.392 | 0.154 | 0.318 | 0.463 | 0.337 | 0.115 | 0.681 | 0.709 | 0.232 | 0.055 |
|  |  |  | 6.0 | 0.050 | 0.134 | 0.228 | 0.391 | 0.154 | 0.323 | 0.462 | 0.337 | 0.115 | 0.684 | 0.708 | 0.233 | 0.056 |
|  |  |  | 5.0 | 0.050 | 0.133 | 0.228 | 0.392 | 0.154 | 0.317 | 0.461 | 0.337 | 0.115 | 0.681 | 0.708 | 0.232 | 0.055 |
|  |  |  | 0 (PIC) | **0.070** | *0.124* | *0.226* | *0.418* | **0.175** | *0.292* | *0.461* | *0.362* | **0.132** | *0.623* | *0.706* | *0.252* | **0.066** |
|  | 3.0 | 7.0 | 3.0 | 0.049 | 0.131 | 0.218 | 0.393 | 0.155 | 0.303 | 0.442 | 0.343 | 0.121 | 0.636 | 0.676 | 0.249 | 0.067 |
|  |  |  | 7.0 | 0.049 | 0.127 | 0.217 | 0.392 | 0.155 | 0.299 | 0.440 | 0.345 | 0.123 | 0.625 | 0.674 | 0.249 | 0.068 |
|  |  |  | 5.0 | 0.050 | 0.128 | 0.218 | 0.392 | 0.155 | 0.299 | 0.443 | 0.341 | 0.120 | 0.625 | 0.674 | 0.249 | 0.068 |
|  |  |  | 0 (PIC) | **0.068** | *0.125* | *0.219* | *0.417* | **0.175** | *0.281* | *0.444* | *0.363* | **0.135** | *0.596* | *0.684* | *0.259* | **0.071** |
| Brawand Tree (n=10) | 4.5 | 5.5 | 4.5 | 0.050 | 0.159 | 0.234 | 0.337 | 0.114 | 0.416 | 0.476 | 0.284 | 0.081 | 0.823 | 0.725 | 0.188 | 0.036 |
|  |  |  | 5.5 | 0.050 | 0.159 | 0.236 | 0.336 | 0.113 | 0.419 | 0.476 | 0.284 | 0.082 | 0.822 | 0.725 | 0.188 | 0.036 |
|  |  |  | 5.0 | 0.050 | 0.162 | 0.234 | 0.338 | 0.114 | 0.418 | 0.474 | 0.285 | 0.082 | 0.825 | 0.726 | 0.186 | 0.035 |
|  |  |  | 0 (PIC) | **0.074** | *0.150* | *0.234* | *0.367* | **0.135** | *0.368* | *0.471* | *0.311* | **0.098** | *0.758* | *0.722* | *0.207* | **0.044** |
|  | 4.0 | 6.0 | 4.0 | 0.049 | 0.157 | 0.233 | 0.337 | 0.114 | 0.414 | 0.472 | 0.288 | 0.084 | 0.813 | 0.720 | 0.190 | 0.037 |
|  |  |  | 6.0 | 0.050 | 0.159 | 0.234 | 0.336 | 0.113 | 0.411 | 0.471 | 0.286 | 0.082 | 0.813 | 0.719 | 0.190 | 0.037 |
|  |  |  | 5.0 | 0.049 | 0.161 | 0.233 | 0.337 | 0.114 | 0.413 | 0.470 | 0.286 | 0.083 | 0.819 | 0.721 | 0.188 | 0.036 |
|  |  |  | 0 (PIC) | **0.074** | *0.148* | *0.233* | *0.368* | **0.136** | *0.366* | *0.471* | *0.310* | **0.097** | *0.752* | *0.719* | *0.209* | **0.045** |
|  | 3.0 | 7.0 | 3.0 | 0.050 | 0.158 | 0.229 | 0.338 | 0.115 | 0.399 | 0.461 | 0.289 | 0.085 | 0.789 | 0.702 | 0.196 | 0.041 |
|  |  |  | 7.0 | 0.049 | 0.150 | 0.224 | 0.339 | 0.115 | 0.388 | 0.456 | 0.292 | 0.087 | 0.774 | 0.696 | 0.201 | 0.043 |
|  |  |  | 5.0 | 0.049 | 0.156 | 0.227 | 0.335 | 0.113 | 0.398 | 0.458 | 0.288 | 0.085 | 0.790 | 0.699 | 0.197 | 0.041 |
|  |  |  | 0 (PIC) | **0.071** | *0.147* | *0.228* | *0.364* | **0.133** | *0.366* | *0.465* | *0.310* | **0.097** | *0.746* | *0.710* | *0.209* | **0.045** |
| Acer Tree (n=55) | 4.5 | 5.5 | 4.5 | 0.049 | 0.576 | 0.247 | 0.129 | 0.017 | 0.989 | 0.496 | 0.105 | 0.011 | 1.000 | 0.745 | 0.062 | 0.004 |
|  |  |  | 5.5 | 0.049 | 0.578 | 0.247 | 0.129 | 0.017 | 0.989 | 0.495 | 0.104 | 0.011 | 1.000 | 0.744 | 0.062 | 0.004 |
|  |  |  | 5.0 | 0.050 | 0.577 | 0.248 | 0.129 | 0.017 | 0.989 | 0.495 | 0.104 | 0.011 | 1.000 | 0.745 | 0.062 | 0.004 |
|  |  |  | 0 (PIC) | **0.080** | *0.503* | *0.247* | *0.144* | **0.021** | *0.971* | *0.495* | *0.117* | **0.014** | *1.000* | *0.745* | *0.070* | **0.005** |
|  | 4.0 | 6.0 | 4.0 | 0.051 | 0.566 | 0.245 | 0.130 | 0.017 | 0.988 | 0.491 | 0.105 | 0.011 | 1.000 | 0.739 | 0.063 | 0.004 |
|  |  |  | 6.0 | 0.050 | 0.570 | 0.245 | 0.129 | 0.017 | 0.988 | 0.491 | 0.105 | 0.011 | 1.000 | 0.738 | 0.064 | 0.004 |
|  |  |  | 5.0 | 0.049 | 0.572 | 0.245 | 0.129 | 0.017 | 0.988 | 0.491 | 0.105 | 0.011 | 1.000 | 0.739 | 0.063 | 0.004 |
|  |  |  | 0 (PIC) | **0.081** | *0.493* | *0.245* | *0.144* | **0.021** | *0.969* | *0.492* | *0.117* | **0.014** | *1.000* | *0.742* | *0.070* | **0.005** |
|  | 3.0 | 7.0 | 3.0 | 0.049 | 0.552 | 0.239 | 0.130 | 0.017 | 0.984 | 0.478 | 0.106 | 0.012 | 1.000 | 0.718 | 0.066 | 0.005 |
|  |  |  | 7.0 | 0.050 | 0.546 | 0.235 | 0.130 | 0.017 | 0.982 | 0.471 | 0.107 | 0.012 | 1.000 | 0.709 | 0.069 | 0.007 |
|  |  |  | 5.0 | 0.048 | 0.552 | 0.237 | 0.129 | 0.017 | 0.984 | 0.475 | 0.106 | 0.012 | 1.000 | 0.713 | 0.067 | 0.006 |
|  |  |  | 0 (PIC) | **0.075** | *0.494* | *0.241* | *0.143* | **0.021** | *0.968* | *0.483* | *0.117* | **0.014** | *1.000* | *0.727* | *0.072* | **0.006** |

## Discussion

### Connection with Maximum Likelihood and GLS Method

Similar to Felsenstein’s PIC method, our OU-PIC method provides a tool for studying correlated evolution under the OU model. Our simulation results showed that appropriate type I error rates and the highest statistical power were achieved when the correct evolution model is specified. The statistical performance of contrast methods drops as the model assumptions deviate from the underlying evolution model. In this section, we discuss the connection between the results of the OU-PIC method and other methods in evaluating trait evolution.

### Connection with the Maximum Likelihood Estimation of the Optimal Trait Value

Under the OU model of trait evolution, the vector of trait values ***X*** = (*X*_1_, *X*_2_,… , *X*_*n*_)^*T*^ follows a multivariate normal distribution, 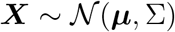. It is straight forward to find the maximum likelihood estimation of *μ* by taking derivatives of the log-likelihood,

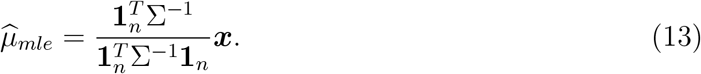

Here **1**_*n*_ is a column vector of *n*-ones. 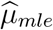 is an unbiased estimator of *μ*.

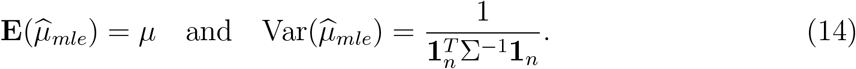

We found that the last intermediate value returned by our algorithm, *w*_*n*-1_, is proportional to 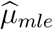, since their coefficients are linearly dependent (Appendix IV). Put in vector forms, we have

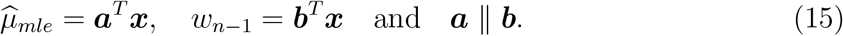

In fact, both of them are independent to all *n* − 1 contrasts (Appendix IV).

#### Connection with the GLS Method for Correlated Evolution

In the GLS regression model, trait *Y* is considered to be determined by trait *X* plus some unspecified noise.

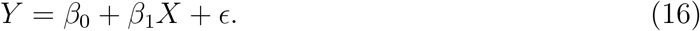

The noise terms *ϵ_i_* are assumed to be Gaussian with zero mean and covariance Σ, *ϵ* ~ *N*(0, Σ). Here, the structure of the covariance matrix describes the phylogenetic relationship among species, and is determined by the phylogenetic tree plus the evolutionary model (Hansen and Martins, 1996). Statistical tests were performed on the slop parameter, *β_1_* in Equation (16) to test hypothesis about correlated evolution. It has long been noticed that when a BM model is assumed, the GLS estimation of the slope parameter in Equation (16), 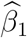, is equivalent to the slope parameter of the ordinary least squares regression of PICs conducted through the origin (Equation 17), 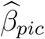

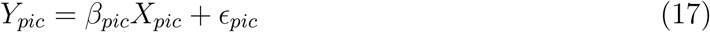

Note that 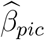 is different from the Pearson correlation coefficient (*r*_*pic*_) of PICs, which is instead related to the slope parameter of the ordinary least squares regression between PICs with an intercept. Blomberg et al. (2012) provided a detailed proof of this equivalence using recursion, but as the authors pointed out, the proof involves some tedious algebraic derivations. We found that this equivalence still holds for our OU-PIC and the GLS regression method under the OU model provided that *κ*_*x*_ = *κ*_*y*_.

We presented a convenient proof for this equivalence by analyzing the properties of the contrast coefficient matrix (Appendix V). In brief, we consider the vector of *n* − 1 contrasts and 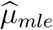, with its coefficient matrix *G*.

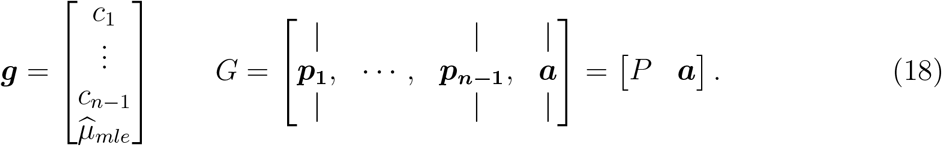

Knowing that elements in ***g*** are normal distributed and independent to each other, *G* is a linear transformation such that *G*^*T*^Σ*G* is diagonalized.

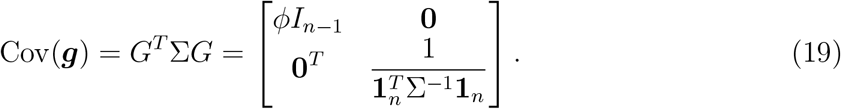

By substituting *G* with *P* and a in equation (19), we arrived at an identity between Σ^-1^ and the coefficient matrix *P*. With that, it becomes straight forward to show that the two regression coefficients 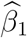 and 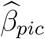 are equivalent.

As discussed previously (Blomberg et al., 2012), this equivalence suggests that ordinary regression analyses using PICs or OU-PICs have the same mathematical shortcomings as GLS methods. However, the algorithm for OU-PIC avoids the explicit calculation of the inverse of matrix Σ and thus would be preferable when the species tree is large.

### Remarks on Method Limitations, Strengths, and Applications

In this study, we investigated the evaluation of correlated evolution when the underlying evolutionary process is Ornstein-Uhlenbeck. By parameterizing the level of correlated evolution under the OU model, problems regarding correlated evolution were formulated to statistical tests or inferences on the correlated evolution parameter from correlated observations (species). Studies on statistical properties of methods for estimating correlation coefficients between two variables could be naturally applied here. Specifically, our OU-PIC algorithm provides a method to decorrelate samples and returns *n* − 1 independent contrasts. We encountered theoretical difficulties when *κ*_*x*_ ≠ *κ*_*y*_though this problem is not unique to the OU-PIC method. According to the regression model of the GLS method, trait *X* is treated as the explanatory variable and trait *Y* is considered as the dependent variable. When *κ*_*x*_ ≠ *κ*_*y*_, only the covariance structure of *Y* is incorporated in the GLS regression model. However, we expect the covariance of the error terms to describe the covariance due to the stochastic evolution of both traits. For this reason, the GLS estimations are also subject to impacted statistical performance. To investigate this problem, we performed simulations to find that OU-PICs still achieve relatively good statistical performance when *κ*_*x*_ and *κ*_*y*_ diverges from each other but remains on the same scale.

Our results showed that the OU-PIC method is the most advantageous when moderate *κ* is expected. If *κ* → 0, the coefficients of OU-PICs get close to the coefficients of PICs. In this case, PICs also achieve good statistical performance in studying correlated evolution. On the contrary, if *κ* is large (*κ* > 10 for species trees we analyzed), the correlation among species resulted from overlapping evolutionary histories is negligible. The statistical performance of PIC may drop dramatically. Instead, raw observations from species could be directly supplied in statistical analyses. Our method has various applications when the trait evolution could be described by an OU process, for example the level of gene expressions that were frequently used in evolutionary studies recently (Rohlfs et al., 2014; Stewart et al., 2019; Kshitiz et al., 2019; Gu et al., 2019). Various studies have shown that the OU model is preferred in describing the evolution of gene expression in many clades (Bedford and Hartl, 2009; Metzger et al., 2017; Yang et al., 2017). Correlated evolution in gene expression has been widely observed among genes as well as between cell types (Ghanbarian and Hurst, 2015; Liang et al., 2018). Our OU-PIC method provides a convenient and statistically sound method to study correlated evolution under an OU process framework.

## Conclusions

In this study, we investigated correlated evolution under the OU model of trait evolution. We derived an extension of Felsenstein’s PIC method under the OU model, called the OU-PIC method. We showed by simulation that our OU-PIC method improves the statistical performance of PIC in studying correlated evolution when stabilizing selection plays a non-negligible role. Our OU-PIC method avoids the calculation of the inverse of the covariance matrix and is straight forward to implement. It has many applications in evolutionary studies including molecular traits such as gene expressions.

## Code Availability

The OU-PIC method described in this manuscript will be available via a python package *oupic* on github. A developing version is also provided as supplementary material for review.

## Acknowledgements

We thank Donghai Pan for helpful discussions. This work was supported by the National Natural Science Foundation of China grant no. 12001401.

## Supplementary Material

Supplementary figures and appendices are available from the Dryad Digital Repository: http://dx.doi.org/10.5061/dryad.xpnvx0kd4.

## Appendix I

We show that under the OU model with the stationary assumption, the covariance of trait *X* between nodes *i* and *j* is:

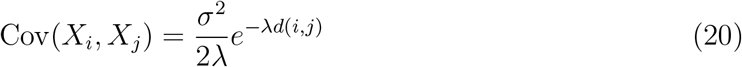

### Proof.

Let *t*_*i*_, *t*_*j*_ be the evolution time between *i*, *j* and their last common ancestor *k*, respectively. Consider the evolutionary process from *k* (*t* = 0) to *i* (*t* = *t*_*i*_),

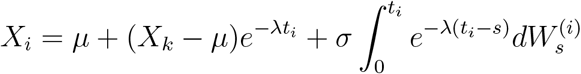

and the process from *k* (*t* = 0) to *j* (*t* = *t*_*j*_),

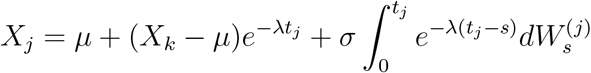

Subscripts for *μ* and λ are omitted here for the simplicity of the notation. With the stationary assumption, 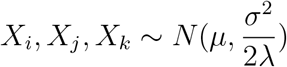. The evolutionary processes are considered independent after the speciation event, i.e. 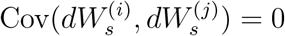. We have,

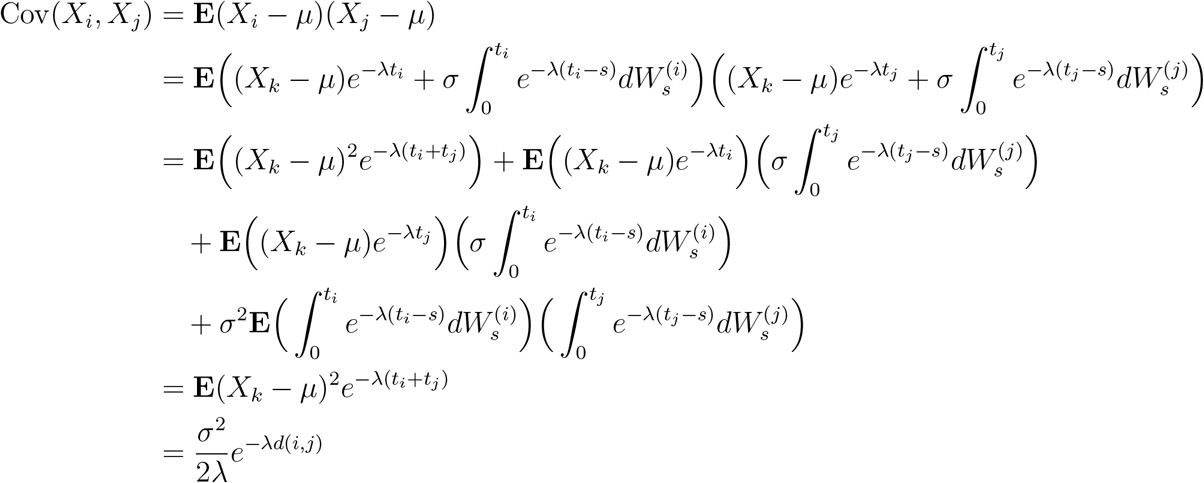

## Appendix II

We show that under the OU model with the stationary assumption, the covariance between two traits *X* and *Y* on node *k* is

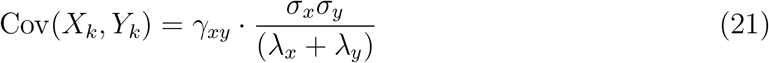

*Proof*. Consider the the evolutionary process from an ancient ancestor *α* (*t* = 0) to node *k* (*t* = *τ*_*k*_). Here *α* is much more ancient than the last common ancestor of the species in the given phylogeny. We have,

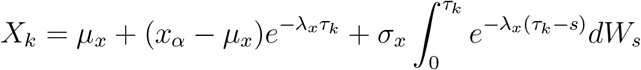

and

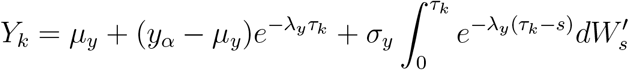

With direct calculation,

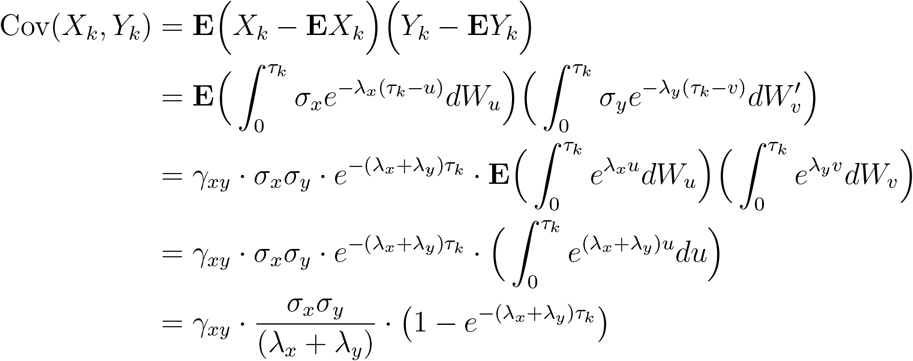

The stationary assumption suggests that the time from the ancient ancestor to the root of phylogeny is long enough and thus can be considered as *τ*_*k*_ → ∞

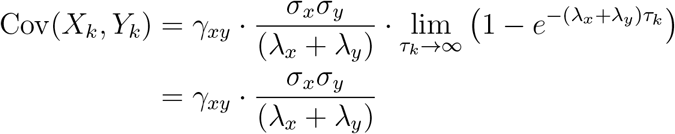

We show that under the OU model with the stationary assumption, the covariance between trait *X* on node *i* and trait *Y* on node *j* is:

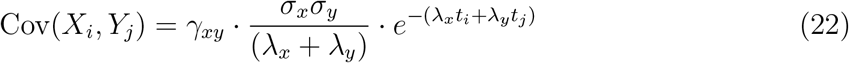

With *t_i_*, *t_j_* as the evolution time between *i*, *j* and their last common ancestor *k*, respectively.

*Proof*. Consider the evolutionary process from *k* (*t* = 0) to *i* (*t* = *t*_*i*_) and the process from *k* (*t* = 0) to *j* (*t* = *t*_*j*_). With the stationary assumption, 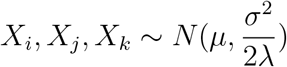. We have,

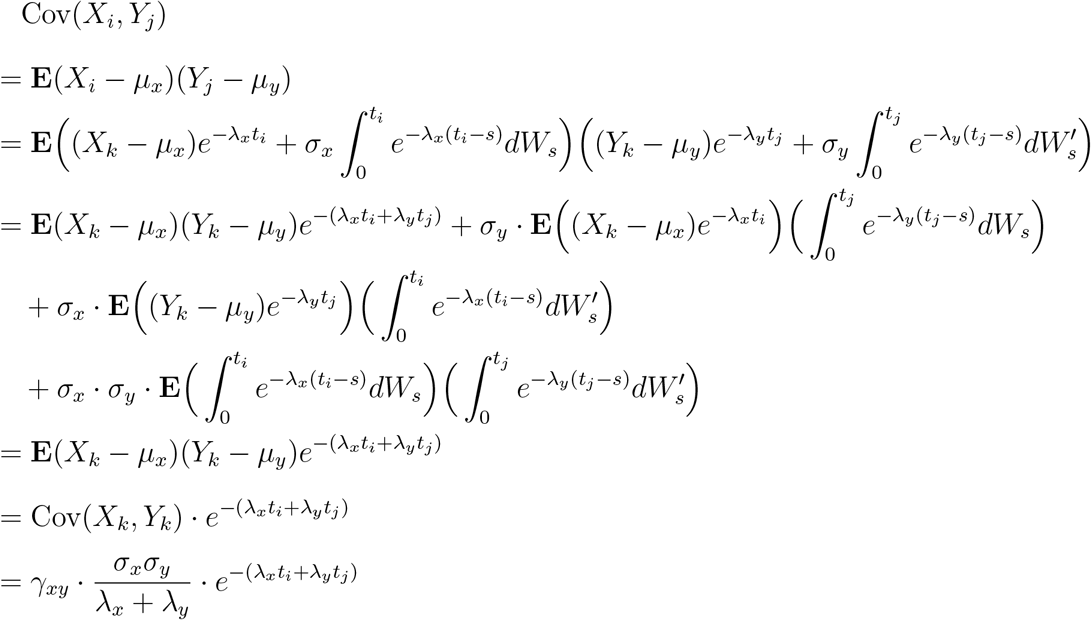

## Appendix III

The coefficients for calculating contrasts and intermediate values in the OU-PIC algorithm were given by the following equations.

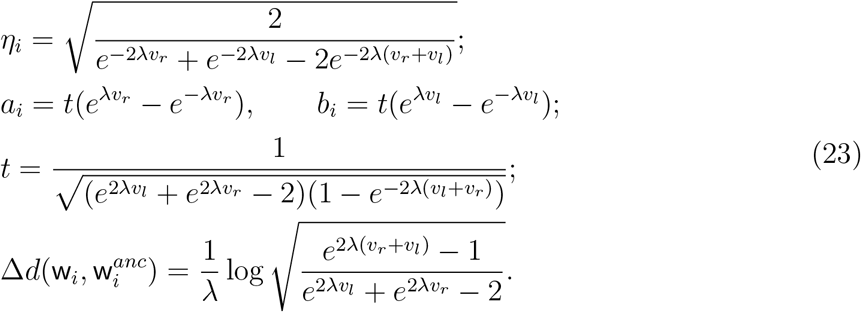

We show that under the OU model with the stationary assumption, contrasts returned by the OU-PIC algorithm have the following properties:

1. *C*_*i*_ ~ *N*(0, *ϕ*), *ϕ* is the normalized variance for all contrasts;
2. *C_i_* are independent to each other;
3. if two traits *X* and *Y* have the same level of stabilizing selection, the Pearson correlation coefficient between their contrasts is *γ*_*xy*_:

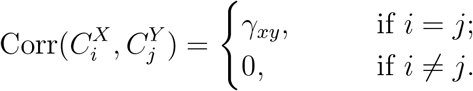

*Proof*. We refer to the phylogeny at the beginning of *i*-th iteration as the *i*-th tree, and denote the divergence time between nodes a and b on the *i*-th tree as *d*^*i*^(a, b). In *i*-th iteration, two leaf nodes x_*l*_ and x_*r*_ were trimmed from the *i*-th tree, and their last common ancestor wi becomes a leaf node in the (*i* + 1)-th tree.

(1) Since *C*_*i*_’s are linear combinations of trait values on leaf nodes, *C*_*i*_’s are also normal distributed. According to the algorithm, variances of *C*_*i*_’s are normalized to 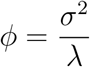. It only remains to show that **E***C*_*i*_ = 0. We show by recursion that 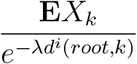 remains a constant for all leaf nodes *k* in all iterations.

(i) When *i* = 1, since the initial species tree has only extant species, the evolution time from the root to any extant species is the same. Denote *d*^1^(root, *k*) = *L*. With the stationary assumption, **E***X*_*k*_ = *μ*. Thus,

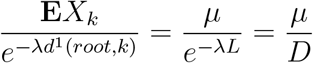
(ii) In *i*-th iteration, **w**_*i*_ becomes a new leaf node with modified branch length, while other nodes remain unchanged. To show that the tree pruning procedure introduced in our algorithm does not change the above constant, we only need to verify that the above ratio for node **w**_*i*_ remains the same in the (*i* + 1)-th tree. The numerator,

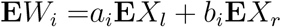 The denominator,

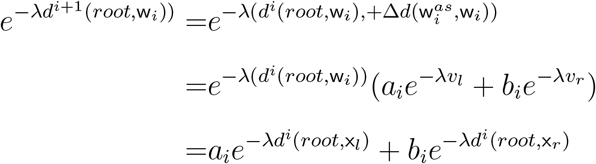 Since 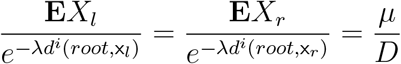, we have,

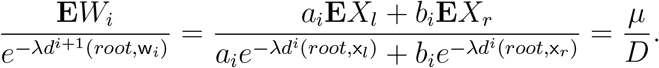 As a result, if 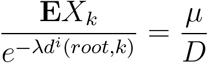 is true for any leaf node *k* in the *i*-th tree, then 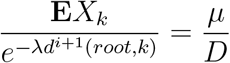 also holds for all leaf nodes in the (*i* + 1)-th tree. With (i) and (ii), 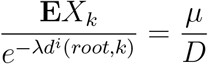 is a constant for all leaf nodes in all iterations. We find the expectation of the *i*-th contrast

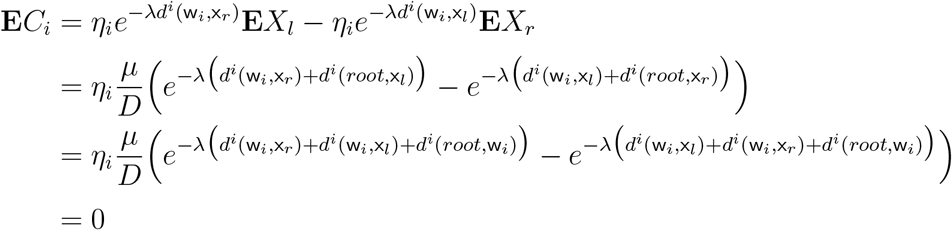
(2) We first find the covariance between node **w**_i_ and any other unmodified leaf node *k* in *i*-th iteration:

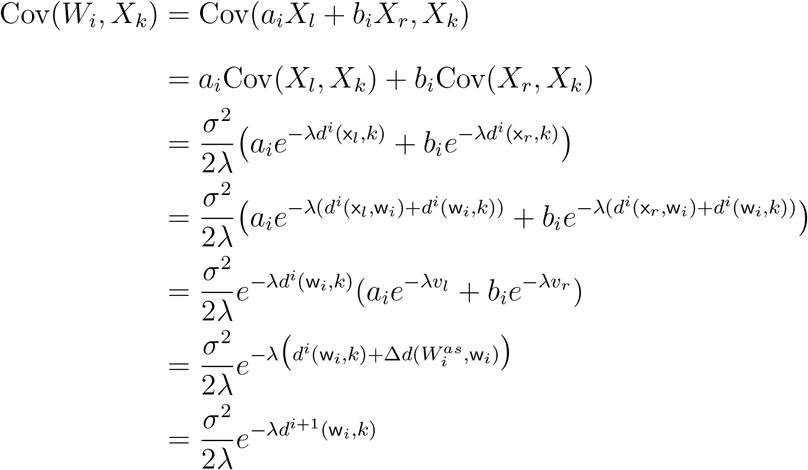 Knowing that 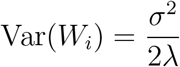, equation (20) about the covariances among leaf nodes still holds in the (*i* + 1)-th tree. Next, we show that the covariance between the contrast *C*_*i*_ and any leaf in the (*i* + 1)-th tree (the pruned tree) is zero. According to the algorithm, Cov(*C*_*i*_, *W*_*i*_) = 0. For other unmodified leaf nodes,

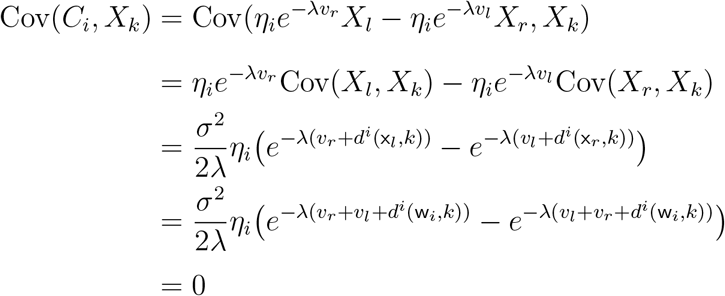 Since any *C*_*j*_ with *j* > *i* is some linear combination of the leaf node values in the *i*-th tree, we have Cov(*C*_*j*_, *C*_*i*_) = 0 for *j* ≠ *i*. As a result, *C*_*i*_ are independent to each other.
(3) Consider two traits *X* and *Y* with λ_*x*_ = λ_*y*_ = λ. According to Equation (23), they obtain the same coefficients for contrasts, intermediate values and branch lengths. First, we show by recursion that equations (21) and (22) hold for all trees when λ_*x*_ = λ_*y*_, i.e.

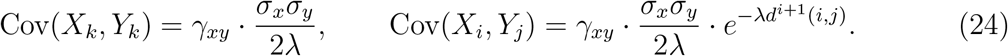 When *i* = 1, we know that equations (24) are true for the original species tree. If equations (24) hold in the *i*-th tree, it is straight forward to verify that the covariance between the values on node **w**_*i*_, 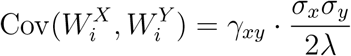, and that the covariance between trait *X* on node **w**_*i*_ and trait *Y* on any unmodified node *k*, 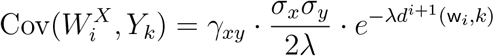 Since the rest of the nodes and branch lengths remains unchanged in *i*-th iteration, equations (24) holds for (*i* + 1)-th tree. As a result, equations (24) are true for trees in all iterations.

Next, we directly verify that 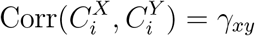

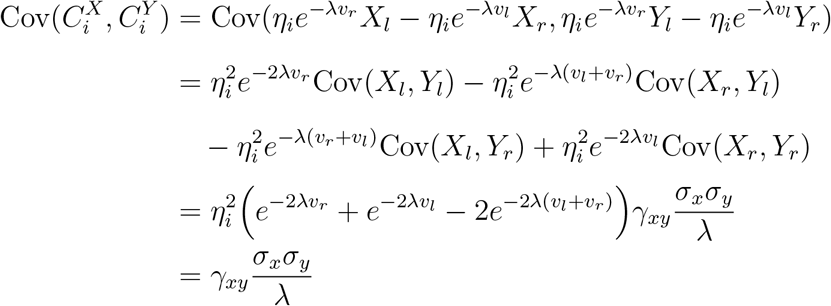

Knowing that 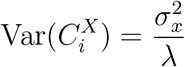 and that 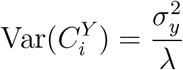, we have

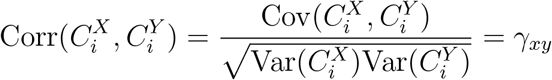

Finally, we show that 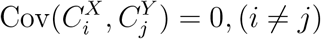. It is sufficient to show this is true when *j* > *i*. When *j* > *i*,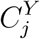 is a linear combination of *Y* values on the (*i* + 1)-th tree. It is sufficient to show that (i) the covariance between 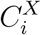 and 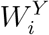 is zero, and that (ii) the covariance between 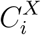 and trait *Y* on other unmodified leaf nodes in *i*-th iteration are all zero.

i. We directly verify the covariance between 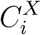 and 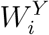

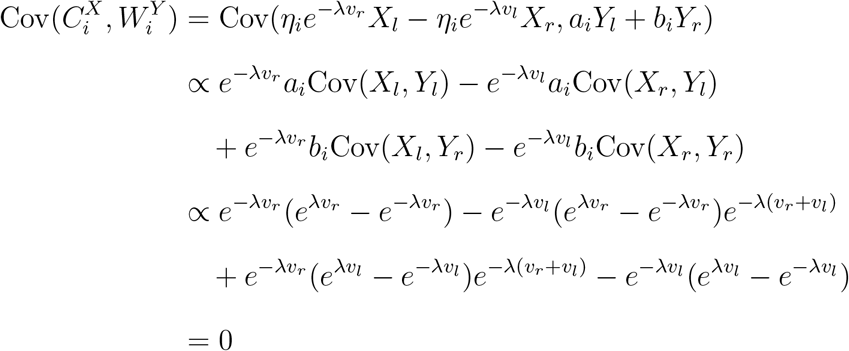
ii. Denote the last common ancestor of node **w**_i_ and node *k* as node **kw**. *k* is a leaf node on *i*-th tree which was not modified in *i*-th iteration.

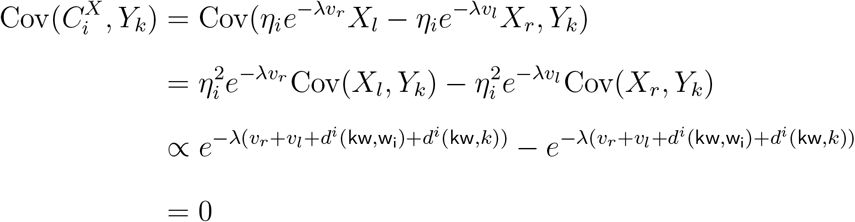

## Appendix IV

We show that *w*_*n*-1_ returned by the OU-PIC algorithm is proportional to the maximum likelihood estimation (MLE) of the optimal value 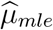.

*Proof.* We first find 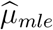, the maximum likelihood estimator of *μ*_*x*_. The vector of trait values ***X*** = (*X*_1_, *X*_2_, … , *X*_*n*_)^*T*^ follows a multivariate normal distribution, 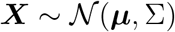. Here, μ = (μ,… , μ)^*T*^ = **1**_*nμ*_. We denote **1**_*n*_ = (1,… , 1)^*T*^, and **0***n* = (0,… , 0)^*T*^.

The probability density of ***X*** is

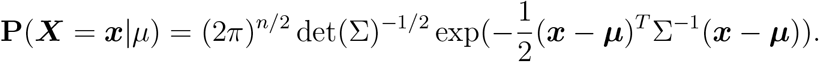

The log-likelihood

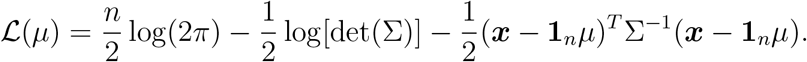

Since 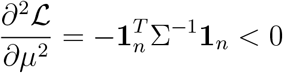. Let 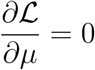 we find the MLE of *μ*,

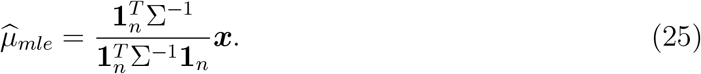

With

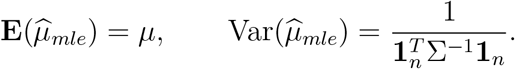

Recall their vector and matrix notations, 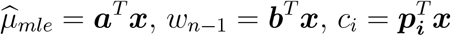 and *P* = (*p*_1_, *p*_2_,… , *p*_*n*-1_). We next show that ***a*** and ***b*** is linearly dependent to each other (***a*** ∥ ***b***).

From equation (25), we have 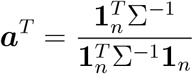. Since 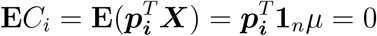, 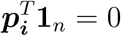, while i ϵ (0, *n* − 1). Thus,

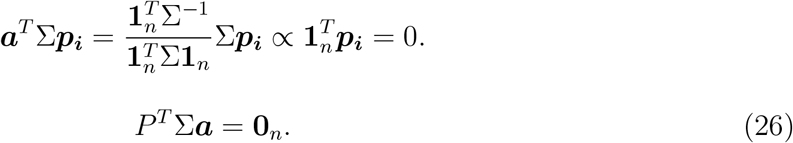

According to Appendix III, the covariance between *C*_*i*_ and all leaf node values on the (*i* + 1)-th tree are zero. Since *W*_*j*_ (*j* > *i*) is a linear combination of leaf node values on the (*i* + 1)-th tree, Cov(*C*_*i*_, *W*_*j*_) = 0, (*j* ≥ *i*). As a result, Cov(*W*_*n*-1_, *C*_*i*_) = 0,*i* ϵ (0,*n* − 1). Thus,

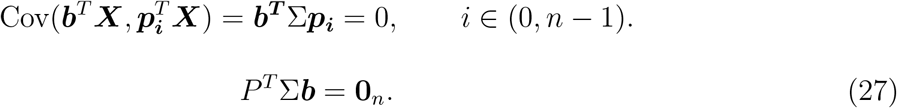

Since *P* is a *n* by *n* − 1 matrix with rank(*P*) = *n* − 1, and rank(Σ) = *n*, *P*^*T*^Σ is a *n* − 1 by *n* matrix with rank(*P*^*T*^Σ) = *n* − 1. With equations (27) and (26), ***a*** and ***b*** are linearly dependent to each other (***a*** ∥ ***b***). *w*_*n*-1_ and 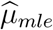 are proportional to each other. The scaling factor between *w*_*n*-1_ and 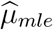 can be identified by normalizing their variance or expectation values.

## Appendix V

We show that the slope parameter of the ordinary least squares regression of OU-PIC conducted through the origin is identical to the slope parameter of generalized least squares regression under a OU model of evolution.

*Proof*. Recall that 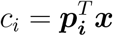 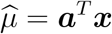and two matrices,

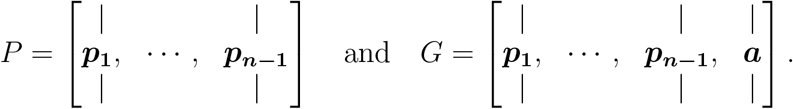

We know that *P*^*T*^Σ*P*. According to Appendix IV, *P*^*T*^Σ***a*** = 0_*n*_, 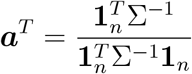 and 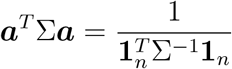. As a result,

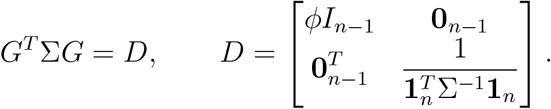

We can derive that

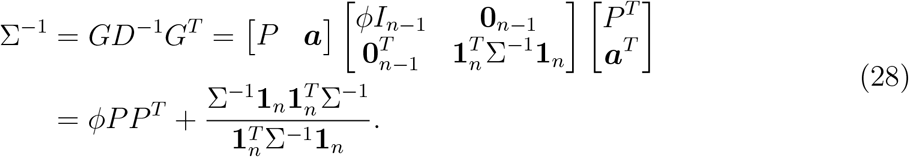

The slope parameter of the ordinary least squares regression of OU-PIC through the origin is 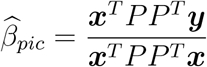

We next find the solution for the generalized least squares problem, *y* = *β*_0_ + *β*_1*x*_, with residual distribution ϵ ~ *N*(0, Σ)

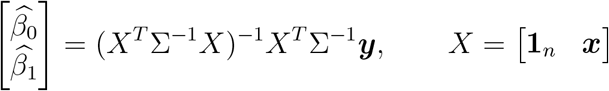

We can solve

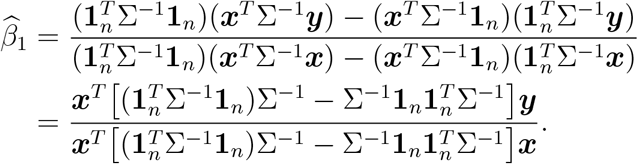

Plug in equation (28) to 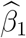, we have

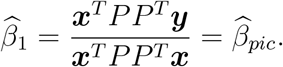

