## Supplementary Figures for "A Phylogenetic Independent Contrast Method under the Ornstein-Uhlenbeck Model and its Applications in Correlated Evolution"

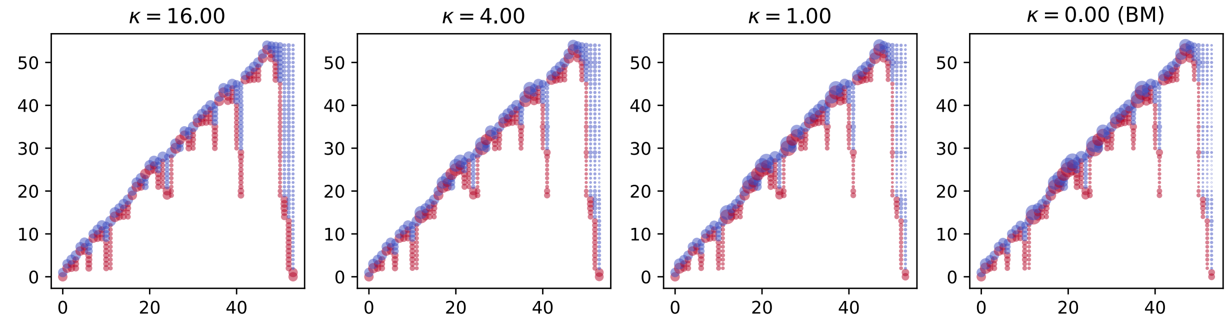


Supplementary Figure 1. Visualization of the coefficient matrix of OU-PICs and PICs for the Acer Tree. The level of relative stabilizing selection $\kappa$ varies under the OU model. When $\kappa = 0$, the evolution model is BM, and the coefficient matrix of PICs is plotted. The horizontal axis in each panel represents the index of contrasts. The vertical axis in each panel represents the index of extant species. To compare coefficients for different $\kappa$, coefficients were normalized according to the coefficients of the first contrast in each panel (the bottom left value). The size of the points is proportional to the absolute value of the normalized coefficients. The color of points represents the sign of coefficients, red for positive, blue for negative.

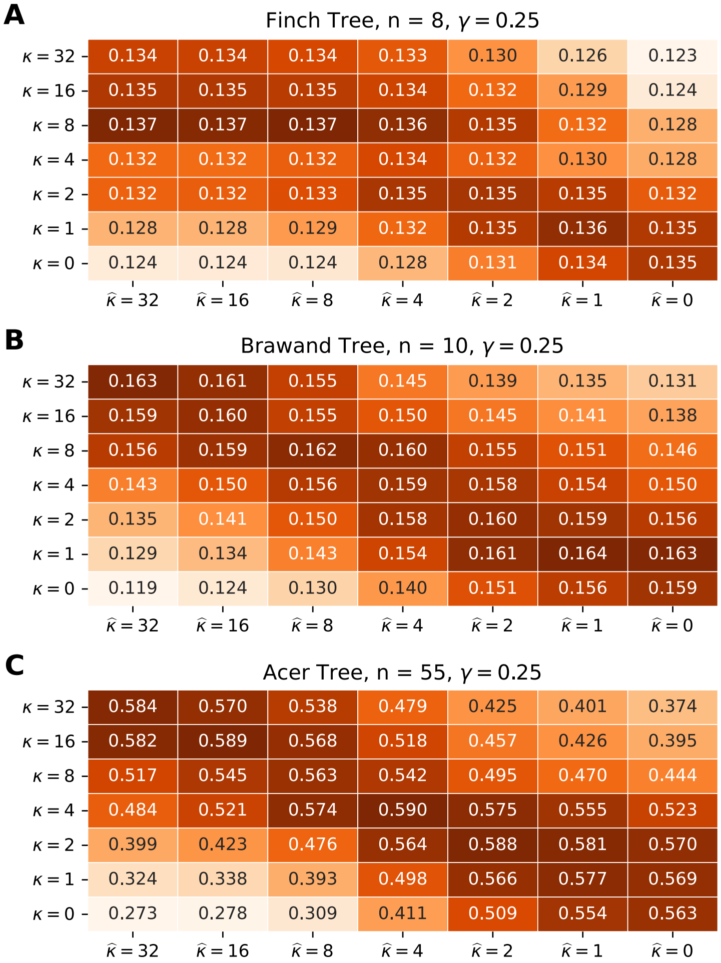


Supplementary Figure 2. The power of test on (A) the Finch Tree, (B) the Brawand Tree, and (C) the Acer Tree given $\gamma = 0.25$. The highest test power is achieved when the correct level of stabilizing selection is specified, i.e. $\widehat{\kappa} = \kappa$.

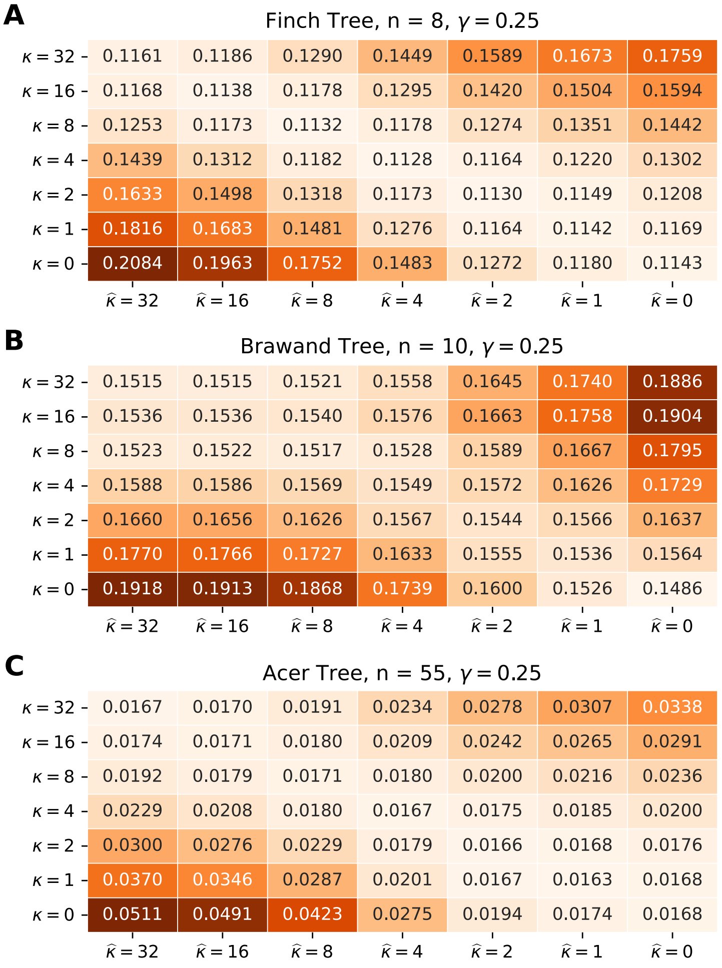


Supplementary Figure 3. The mean squared errors in estimating $\gamma$ on (A) Finch Tree, (B) Brawand Tree, and (C) Acer Tree given $\gamma = 0.25$. The lowest mean squared error is achieved when $\widehat{\kappa} = \kappa$.

**A**

Finch Tree, n = 8, $\gamma = 0.50$

| | $\widehat{\kappa} = 32$ | $\widehat{\kappa} = 16$ | $\widehat{\kappa} = 8$ | $\widehat{\kappa} = 4$ | $\widehat{\kappa} = 2$ | $\widehat{\kappa} = 1$ | $\widehat{\kappa} = 0$ |
|---|---|---|---|---|---|---|---|
| $\kappa = 32$ | 0.0814 | 0.0835 | 0.0920 | 0.1049 | 0.1164 | 0.1234 | 0.1307 |
| $\kappa = 16$ | 0.0842 | 0.0821 | 0.0854 | 0.0948 | 0.1050 | 0.1118 | 0.1191 |
| $\kappa = 8$ | 0.0914 | 0.0846 | 0.0812 | 0.0848 | 0.0921 | 0.0980 | 0.1052 |
| $\kappa = 4$ | 0.1082 | 0.0979 | 0.0875 | 0.0832 | 0.0857 | 0.0899 | 0.0963 |
| $\kappa = 2$ | 0.1219 | 0.1110 | 0.0965 | 0.0851 | 0.0817 | 0.0830 | 0.0876 |
| $\kappa = 1$ | 0.1370 | 0.1257 | 0.1089 | 0.0923 | 0.0833 | 0.0813 | 0.0832 |
| $\kappa = 0$ | 0.1574 | 0.1472 | 0.1297 | 0.1081 | 0.0914 | 0.0842 | 0.0812 |

**B**

Brawand Tree, n = 10, $\gamma = 0.50$

| | $\widehat{\kappa} = 32$ | $\widehat{\kappa} = 16$ | $\widehat{\kappa} = 8$ | $\widehat{\kappa} = 4$ | $\widehat{\kappa} = 2$ | $\widehat{\kappa} = 1$ | $\widehat{\kappa} = 0$ |
|---|---|---|---|---|---|---|---|
| $\kappa = 32$ | 0.1136 | 0.1136 | 0.1140 | 0.1167 | 0.1234 | 0.1307 | 0.1422 |
| $\kappa = 16$ | 0.1144 | 0.1144 | 0.1145 | 0.1170 | 0.1236 | 0.1311 | 0.1430 |
| $\kappa = 8$ | 0.1133 | 0.1133 | 0.1130 | 0.1143 | 0.1196 | 0.1262 | 0.1371 |
| $\kappa = 4$ | 0.1164 | 0.1162 | 0.1147 | 0.1128 | 0.1148 | 0.1195 | 0.1285 |
| $\kappa = 2$ | 0.1259 | 0.1256 | 0.1230 | 0.1174 | 0.1145 | 0.1156 | 0.1209 |
| $\kappa = 1$ | 0.1316 | 0.1313 | 0.1280 | 0.1200 | 0.1136 | 0.1120 | 0.1141 |
| $\kappa = 0$ | 0.1486 | 0.1482 | 0.1445 | 0.1341 | 0.1231 | 0.1174 | 0.1147 |

**C**

Acer Tree, n = 55, $\gamma = 0.50$

| | $\widehat{\kappa} = 32$ | $\widehat{\kappa} = 16$ | $\widehat{\kappa} = 8$ | $\widehat{\kappa} = 4$ | $\widehat{\kappa} = 2$ | $\widehat{\kappa} = 1$ | $\widehat{\kappa} = 0$ |
|---|---|---|---|---|---|---|---|
| $\kappa = 32$ | 0.0107 | 0.0109 | 0.0122 | 0.0151 | 0.0179 | 0.0198 | 0.0219 |
| $\kappa = 16$ | 0.0113 | 0.0111 | 0.0116 | 0.0135 | 0.0157 | 0.0172 | 0.0189 |
| $\kappa = 8$ | 0.0123 | 0.0115 | 0.0110 | 0.0118 | 0.0133 | 0.0145 | 0.0159 |
| $\kappa = 4$ | 0.0149 | 0.0135 | 0.0116 | 0.0107 | 0.0112 | 0.0118 | 0.0128 |
| $\kappa = 2$ | 0.0199 | 0.0183 | 0.0151 | 0.0119 | 0.0111 | 0.0114 | 0.0120 |
| $\kappa = 1$ | 0.0253 | 0.0236 | 0.0195 | 0.0136 | 0.0113 | 0.0110 | 0.0113 |
| $\kappa = 0$ | 0.0342 | 0.0328 | 0.0280 | 0.0180 | 0.0127 | 0.0113 | 0.0110 |

Supplementary Figure 4. The mean squared errors in estimating $\gamma$ on (A) Finch Tree, (B) Brawand Tree, and (C) Acer Tree given $\gamma = 0.5$. The lowest mean squared error is achieved when $\widehat{\kappa} = \kappa$.
